# Time-dependent regulation of cytokine production by RNA binding proteins defines T cell effector function

**DOI:** 10.1101/2021.11.03.467112

**Authors:** Branka Popović, Benoît P. Nicolet, Aurélie Guislain, Sander Engels, Anouk P. Jurgens, Natali Paravinja, Julian J. Freen-van Heeren, Floris P.J. van Alphen, Maartje van den Biggelaar, Fiamma Salerno, Monika C. Wolkers

## Abstract

Potent T cell responses against infections and malignancies depend on the release of effector molecules, such as pro-inflammatory cytokines. Because effector molecules can be toxic, their production is tightly regulated through post-transcriptional events at 3’ Untranslated Regions (3’UTRs). RNA binding proteins (RBPs) were shown to be key regulators herein. With an RNA aptamer-based capture assay from human T cells, we identified >130 RBPs interacting with the *IFNG, TNF* and *IL2* 3’UTRs in human T cells. T cell activation altered RBP-RNA interactions, revealing that RBP-target mRNA interactions rapidly respond to stimulation. Furthermore, we uncovered the intricate and time-dependent regulation of cytokine production by RBPs: whereas HuR supports early cytokine production, ZFP36L1, ATXN2L and ZC3HAV1 dampen and shorten the production duration, each at different time points. Strikingly, even though ZFP36L1 deletion did not phenotypically rescue T cell dysfunction in tumors, the increased production of cytokines and cytotoxic molecules resulted in superior anti-tumoral T cell responses *in vivo*. Our findings thus show that identifying RBP-RNA interactions reveals key modulators of T cell responses in health and disease.

## Introduction

T cells are critical players in our defense against infections and malignancies. Their production of effector molecules such as granzymes and proinflammatory cytokines is key. Interferon gamma (IFN-γ) and Tumor necrosis factor (TNF) are major contributors to anti-microbial and anti-tumoral T cell responses (*1, 2*), with the most potent T cells co-producing the survival-inducing cytokine IL-2 (*3*–*7*). Whereas the activity of cytokines on target cells is well characterized, the molecular switches that dictate their production is not well understood. Recently, post-transcriptional regulation (PTR) was found to dictate the cytokine production levels (*8–10*). We showed that the strength of T cell receptor signaling together with co-stimulation defines the synthesis and degradation rate of cytokine mRNA in T cells, as well as their translation efficiency (*11–13*).

The 3’Untranslated region (3’UTR) of the mRNA is a major contributor in PTR. For instance, germ-line deletion of cis-elements such as AU-rich sequences (AREs) from the *Tnf* and *Ifng* 3’UTR results in hyperinflammation and immunopathology (*14, 15*). Conversely, tumor-infiltrating T cells (TILs) fail to produce IFN-γ protein despite their continuous high expression of *Ifng* mRNA (*16*). Germ-line deletion of AREs from the *Ifng* 3’UTR restored IFN-γ protein production in murine TILs, and boosted their anti-tumoral potency (*16*). Notably, this augmented protein production was conserved in human T cells (*17*).

RNA binding proteins (RBPs) are critical mediators of PTR that define the fate of mRNA (*18*). For instance, ZC3H12A (Regnase-1) and Rc3h1 (Roquin-1) prevent naïve CD4^+^ T cells from exiting quiescence by destabilizing mRNAs encoding regulators of CD4^+^ T cell differentiation and function (*19*). In contrast, Arid5a promotes T cell differentiation by stabilizing *Stat3* and *Ox40* mRNA (*20, 21*). We found that ZFP36L2 blocks translation of pre-formed cytokine mRNAs in resting memory T cells, thereby preventing aberrant protein production from these ready-to-deploy mRNAs (*22*). Importantly, reactivating memory T cells releases the *Tnf* and *Ifng* mRNA from ZFP36L2, thereby licensing the immediate cytokine production of memory T cells (*22*). Thus, RBPs substantially impact the acquisition and execution of T cell effector function. Which RBPs interact with cytokine 3’UTRs upon T cell activation and define their protein production levels is however not well understood. A comprehensive study on RBP interactions with target mRNAs in T cells is lacking.

More than 2000 RBPs have been annotated in mammalian cells (*23–26*). RNA-RBP interaction maps have been generated for 150 RBPs with Cross-linked immunoprecipitation (CLIP) methods (*27*). However, both mRNA expression and RBP expression is cell-type specific (*28*), thereby prohibiting direct translation of the RBP interaction maps from epithelial cell lines to human T cells. Furthermore, CLIP depends on the availability of suitable antibodies or on tagging endogenous RBPs (*29*). To identify the RBPs that interact with cytokine mRNAs in an unbiased manner, an RNA-centric approach is required. With an RNA aptamer pull-down approach from primary human T cell lysates using full-length cytokine 3’UTRs, we present the first comprehensive analysis of RBP-mediated regulation of T cell effector function, and we reveal the potential of RBP modulation in defining T cell responses to target cells.

## Results

### Cytokine 3’UTRs define the protein production in murine and human T cells

To determine how cytokine 3’UTRs contribute to protein production, we retrovirally transduced peripheral blood-derived human T cells with GFP reporter constructs containing the full length 3’UTR of the human *GZMB, IFNG, TNF* or *IL2* mRNA. The empty GFP construct served as control (GFP_control_). Whereas *GZMB* 3’UTR only slightly reduced the GFP expression levels in nonactivated CD8^+^ T cells compared to GFP_control_, the presence of cytokine 3’UTRs conferred a substantial loss of the GFP signal (Fig. 1A, B). T cell activation with PMA/Ionomycin for 6 h and 16 h marginally increased the protein expression for GFP_control_ and GFP-*GZMB-*3’UTR. In contrast, the GFP expression levels for cytokine 3’UTR-containing constructs increased by a 2-to 8-fold (Fig. 1A, C). No loss of regulation was observed for the endogenous cytokine production (SI Appendix, Fig. S1A). However, the increased GFP levels for cytokine 3’UTRs in activated T cells did not reach the levels of GFP_control_, indicating further regulation of cytokine production in activated T cells. Similar results were obtained with human CD4^+^ T cells, and with murine OT-I T cell receptor (TCR) transgenic CD8^+^ T cells expressing the murine *Gzmb, Gzma* and *Prf1* 3’UTR, compared to the *Ifng, Tnf* or *Il2* 3’UTRs (SI Appendix, Fig. S1B-E). The profound regulation of protein expression by cytokine 3’UTRs is thus conserved.

**Fig. 1.**
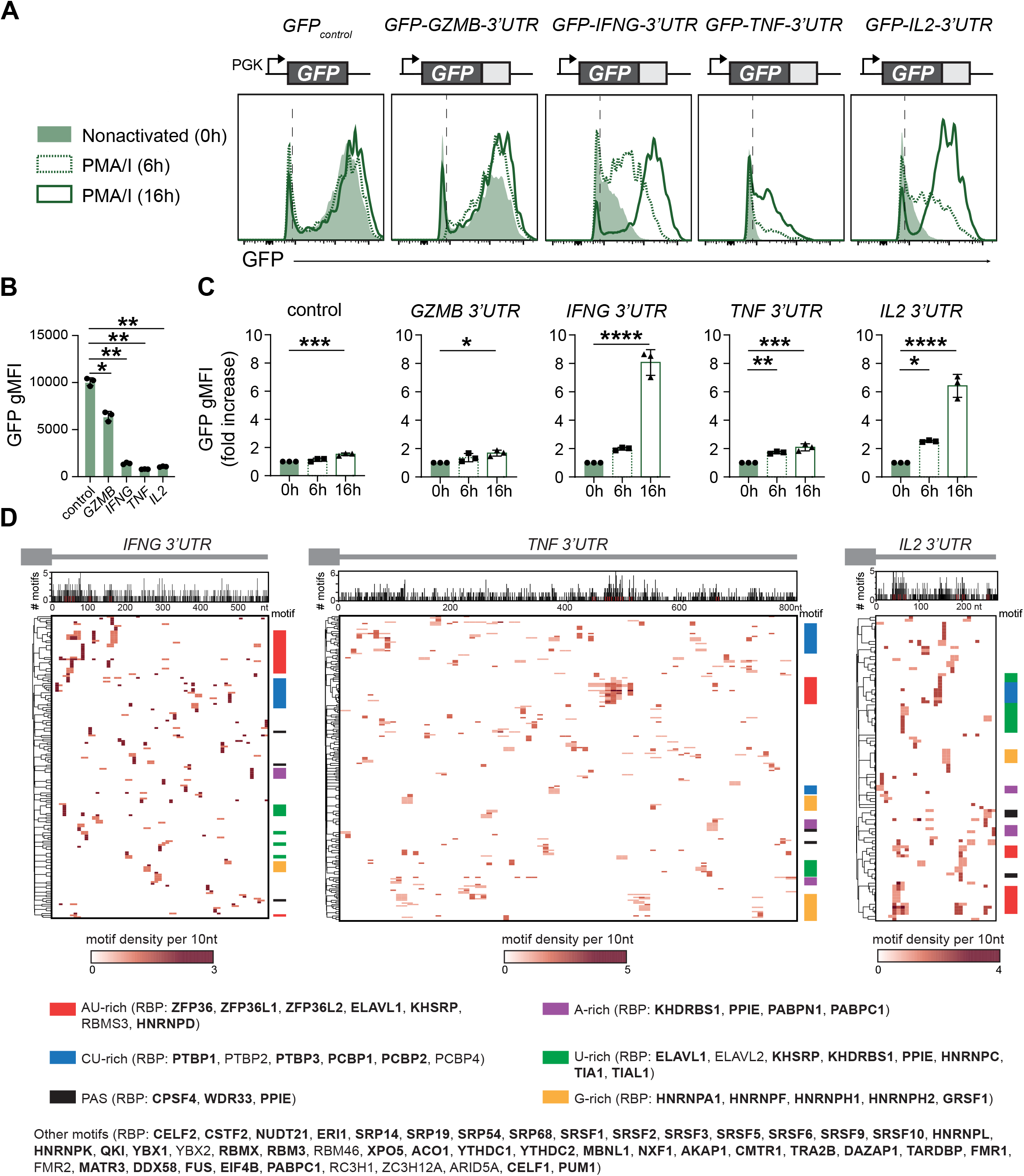
Conserved regulation of protein production by cytokine 3’UTRs. (*A*) Human T cells were transduced with GFP reporter constructs containing indicated human 3’UTRs, or with GFP empty control. Representative GFP expression of nonactivated CD8^+^ T cells (green histograms), or of T cells activated with PMA/Ionomycin (PMA/I) for 6 and 16 h (dashed and solid green lines, respectively). (*B*) GFP gMFI in nonactivated human CD8^+^ T cells. Data depict mean ± s.d. of 3 donors and are representative of at least 2 independently performed experiments. (*C*) Fold increase of GFP gMFI upon activation with PMA/I compared to nonactivated GFP-expressing human CD8^+^ T cells. Data are presented as mean ± s.d. of 3 donors, and are representative of 2 independently performed experiments. (*B-C*) One-way ANOVA with Dunnett’s multiple comparison to the control; *P < 0.05; **P < 0.01; ***P < 0.001; ****P < 0.0001. (*D*) RBP binding motifs in cytokine 3’UTRs. (top) Graphs indicate the number of motifs per 3’UTR sequence. (bottom) Heatmaps represent motif density per 3’UTR sequence. Clusters are color-coded based on the motif sequence: AU-rich, CU-rich, A-rich, U-rich, G-rich and PAS (poly(A) signal). Putative RBP interactors are indicated in brackets, with RBPs detected in whole-cell T cell lysates by mass spectrometry indicated in bold.

We next questioned which RBPs can interact with cytokine 3’UTRs. We assessed putative RBP binding sites *in silico* with the ATtRACT database (*30*), which contains 2297 consensus motifs for >140 RBPs. All three cytokine 3’UTRs contain A-rich, AU-rich, CU-rich, U-rich and G-rich motifs, and several poly(A) sites (PAS), which is also observed for the *GZMB* 3’UTR (Fig. 1D, SI Appendix, Fig. S1F). These motifs are potential RBP binding hubs for ZFP36 family members, ELAVL1, KHSRP, PPIE, CSTF2, NUDT21 and other RBPs such as members of the protein families PCBP, PTBP, HNRNP, SRP, SRSF, PABP, RBM and YTHDC (Fig. 1D, SI Appendix, Fig. S1F). In conclusion, the cytokine 3’UTRs contain a wide range of putative RBP binding sites, which warrants the identification of their actual RBP interactors.

### Identification of RBPs interacting with full length cytokine 3’UTRs

To experimentally identify the RBPs that interact with cytokine 3’UTRs in human T cells, we generated *in vitro* transcribed streptavidin-binding 4xS1m RNA aptamers (*31*) with full length 3’UTRs of *IFNG, TNF*, and *IL2* (Fig. 2A). The empty 4xS1m RNA aptamer served as control. The 4xS1m aptamer system efficiently purifies RNPs, because its improved affinity to streptavidin allows for stringent washes to reduce background, without losing bona fide RBP interactions (*31*). Because RBP expression is cell-type specific (*28*), we performed the RBP pull-down with full cell lysates from human primary CD3^+^ T cells, allowing to interrogate both nuclear and cytosolic RBP-3’UTR interactions. T cells were isolated from 3 pools of 40 donors as RBP source and activated with α-CD3/α-CD28 prior to culture for 5 days in low IL-2 to generate nonactivated effector T cells. At this time point CD69 expression was low and cytokine production was undetectable (SI Appendix, Fig. S2A). Upon capture of RNA-RBP complexes with streptavidin beads, RNA-interacting proteins were identified by mass spectrometry (MS).

**Fig. 2.**
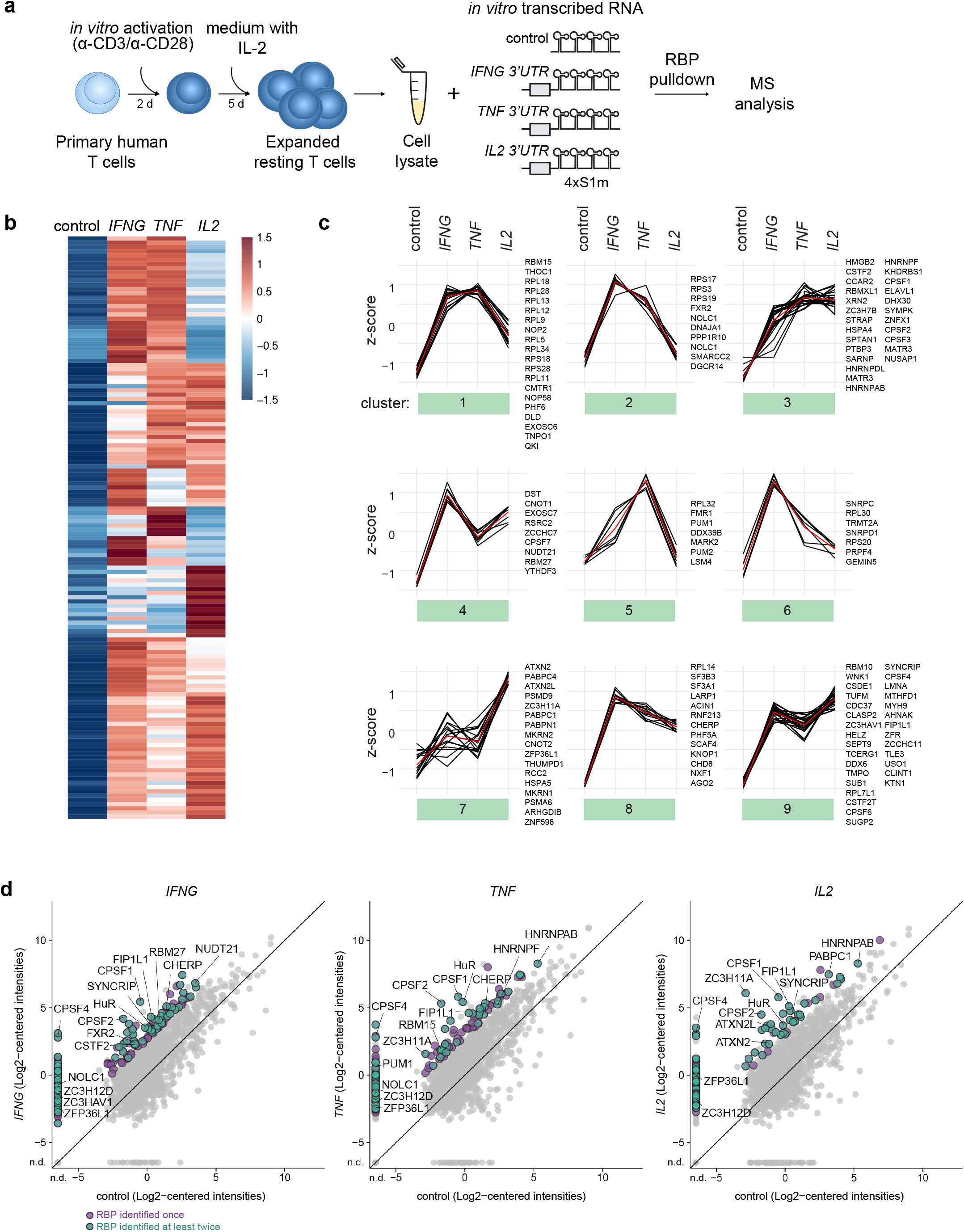
Identification of RNA binding proteins that interact with *IFNG, TNF* and *IL2* 3’UTRs. (*A*) Schematic representation of the RBP pull-down assay. T cell lysates were generated from 40 pooled donors that were activated for 2 days with α-CD3/α-CD28 and rested for 5 days with rhIL-2. Lysates were incubated with *in vitro* transcribed cytokine 3’UTR-containing 4xS1m RNA aptamers, or with empty 4xS1m RNA aptamers (control). Protein binding to 4xS1m RNA aptamers was identified by label-free MS analysis. (*B*) Heatmap of RBPs that were reproducibly enriched upon pull-down with 4xS1m RNA aptamers containing the *IFNG, TNF* or *IL2* 3’UTR, compared to control (n=138). Enrichment cut-off was LFC>4. Color scale of heatmaps represents Z-scored log2 median-centered averaged intensities. Numbers indicate clusters. (*C*) Cluster analysis of RBP interaction specificities using data from (B). Red line depicts the average expression. (*D*) Comparison of protein raw log2 median-centered intensities from the *IFNG* (left panel), *TNF* (middle panel) and *IL2* (right panel) pull-down versus control. Expression levels of duplicates (*IFNG*) or triplicates (*TNF* and *IL2*) were averaged. Purple dots depict RBP candidates identified in one pull-down experiment with an LFC>4 compared to control. Green dots denote RBP candidates reproducibly identified in at least 2 independently performed pull-down experiments, each with T cells pooled from 40 donors. Gray dots depict all proteins identified in representative experiment. n.d.= not detected.

We detected in total 1808 proteins, of which 598 proteins were detected in at least 2 of the 3 replicates with a log_2_ fold change (LFC)>4 in the cytokine 3’UTR aptamer pull-downs compared to the empty aptamer control (SI Appendix, Fig. S2B-D, Dataset S1). Of these, 307 proteins (51.3%) were experimentally confirmed or computationally predicted RBPs (n=222 and n=85, respectively; *see Supplementary methods*). In the downstream analysis, we focused on RBPs that were enriched in at least two independently performed experiments. This included 138 RBPs of which 93 RBPs were enriched for the *IFNG* 3′ UTR, 69 RBPs for *TNF* 3′ UTR, and 82 RBPs for the *IL2* 3′ UTR (Fig. 2B, D, Dataset S1). Gene ontology (GO) analysis on all 138 detected RBPs revealed that the terms ‘RNA processing’ and ‘(m)RNA metabolic processes’ were enriched for all three 3’UTRs (SI Appendix, Fig. S2E). ‘Translation’ was in particular enriched in *IFNG-* and *TNF*-associated RBPs, and ‘RNA splicing’ in *IFNG* and *IL2-*associated RBPs (SI Appendix, Fig. S2E, Dataset S2).

Some RBPs were specifically enriched for one cytokine 3’UTR, such as Fragile X mental retardation syndrome-related protein 2, FXR2, and RBM27 for *IFNG* 3’UTR, stress granule-components ATXN2 and ATXN2L and poly(A)-tail-binding proteins PABPC1 for *IL2* 3’UTR, and Pumilio homolog 1, PUM1, for *TNF* 3’UTR (Fig. 2B-D and Dataset S1). Other RBPs engaged with more than one cytokine 3’UTR, including NOLC1, THOC1 and CHERP (*IFNG* and *TNF*), the splicing factor SYNCRIP (*IFNG* and *IL2*), and HNRNPAB (*TNF* and *IL2*; Fig. 2B-D and Dataset S1). RBPs interacting with all three cytokine 3’UTRs include components of cleavage and polyadenylation specificity factor complex, CPSF4, FIP1L1, the zinc-finger protein ZC3HAV1 and U-binding protein HuR (ELAVL1) (Fig. 2B-D and Dataset S1). ZFP36L2, which we previously identified to bind to preformed *Ifng* and *Tnf* mRNA in resting memory T cells (*22*), was also detected, but with LFCs of 2.5 (*IFNG*), 1.9 (*TNF*) and 3.3 (*IL2*) did not meet our stringent cut off of LFC>4 (Dataset S1). Notably, several RBPs we detected interacting with cytokine 3’UTRs were also found in the RBPome data set from Hoefig et al (*26*), and included SYNCRIP, TIA-1, CSDE1, HuR, PUM2, and PTBP3. In conclusion, using primary human T cell lysates as bait, we identified 138 RBPs that display specific or preferential interaction profiles with full length cytokine 3’UTRs.

### The RBP binding landscape alters upon T cell activation

We and others previously showed that post-transcriptional events are critical to control cytokine production in T cells. External triggers can alter the expression levels of RBPs and/or result in post-translational modifications, which in turn can alter their RNA binding capacity (*22, 32, 33*). To determine how T cell activation modulates the RBP binding landscape, we performed the aptamer pull-down with lysates from 2 h PMA/Ionomycin-activated CD3^+^ T cells, a setup that allows for rapid and homogenous T cell activation (Fig. 3A). We detected in total 1596 proteins (SI Appendix, Fig. S2F, Dataset S3), of which 443 proteins were enriched with LFC>4 in cytokine 3’UTR samples compared to empty aptamer control. 244 proteins (55.1%) were annotated as RBPs, and 77 RBPs were detected in two independently performed experiments (Fig. 3A, Dataset S3). Using the cytokines 3’UTRs as controls for each other, we found that 58 of these 77 RBPs displayed enriched binding to *IFNG* 3′UTR, 63 RBPs to *TNF* 3′UTR, and 22 RBPs to the *IL2* 3′UTR (Fig. 3A). Because the number of reproducibly enriched RBPs with an LFC>4 for IL2 3’UTR was low, we excluded this data set from GO term enrichment analysis. RBPs interacting with the *IFNG* and *TNF* 3’UTR in activated T cells included ‘RNA processing’ and ‘(m)RNA metabolic processes’ (SI Appendix, Fig. S2G, Dataset S4).

**Fig. 3.**
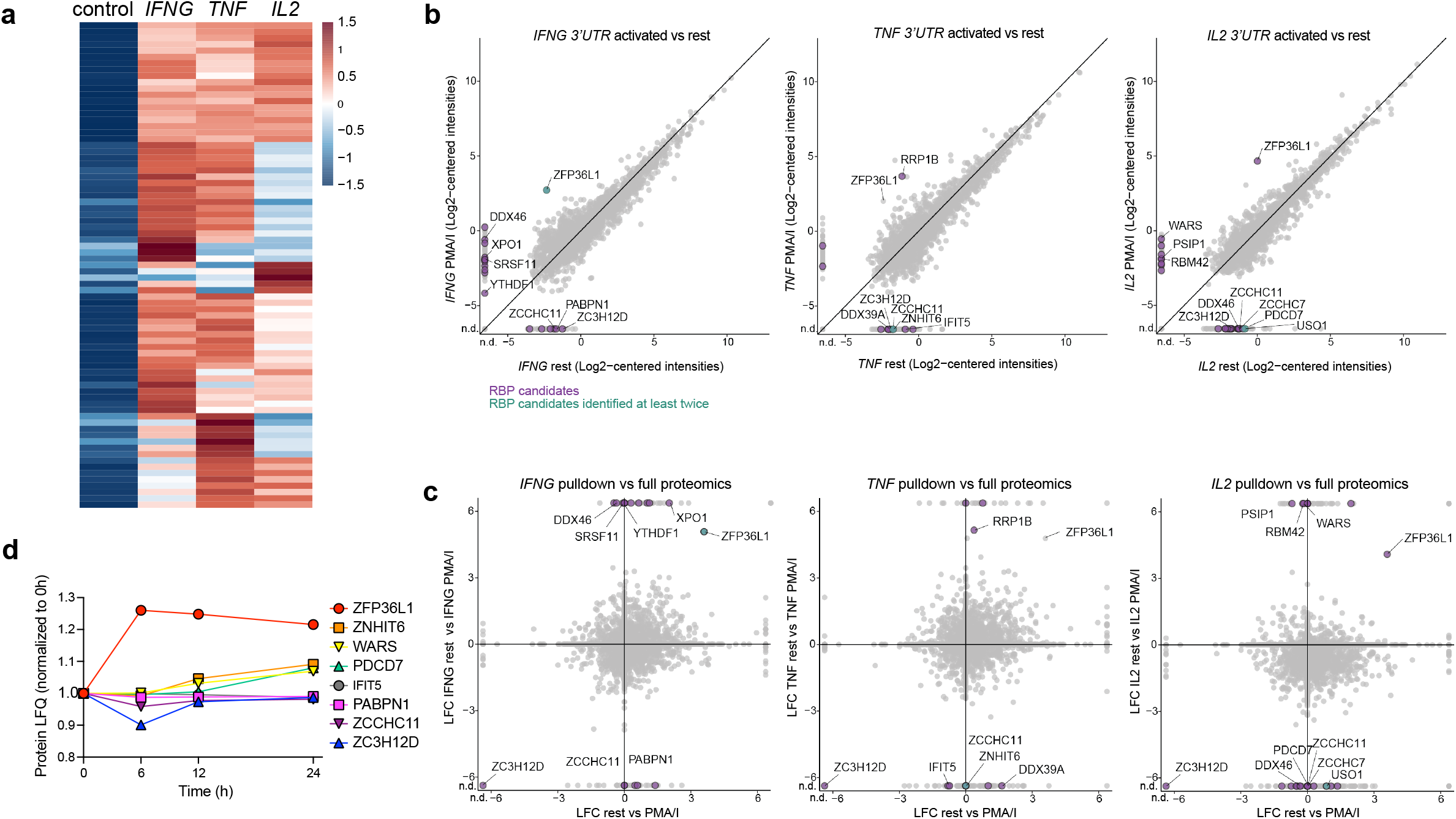
The RBP landscape of *IFNG, TNF* and *IL2* 3’UTR alters upon T cell activation. (*A*) Heatmap of reproducibly enriched RBPs from cytosolic lysates of PMA/I-activated T cells (2 h) upon pull-down with cytokine 3’UTR-containing 4xS1m aptamers compared to empty 4xS1m aptamers (control) (n=77). Enrichment cut-off: LFC>4. Color scale of heatmaps represents Z-scored log2 median-centered averaged intensities. (*B*) Comparison of protein raw log2 median-centered intensities from *IFNG* (left panel), *TNF* (middle panel) and *IL2* (right panel) 3’UTR pull-downs from nonactivated T cells versus activated T cells. (*C*) Log2 Fold Change (LFC) of log2 intensities from *IFNG* (left panel), *TNF* (middle panel) and *IL2* (right panel) 3’UTR pull-down between nonactivated and activated T cells and LFC of protein log2 intensities from the whole-cell proteome of the cytosolic lysates between nonactivated and activated T cells. (*B-C*) Expression levels of triplicates were averaged. Green and purple dots show proteins with enrichment LFC>4 compared to control and LFC<-4 or >4 between nonactivated and activated T cells. Purple dots depict RBP candidates identified in one pull-down experiment, green dots denote RBP candidates reproducibly identified in at least 2 independently performed pull-down experiments, each with T cells pooled from 40 donors. Gray dots depict all proteins identified in one experiment. n.d.= not detected. (D) Expression levels of indicated RBPs in human memory T cells after activation for 0, 6, 12 and 24 h (*34*).

We next studied whether RBPs alter their RNA binding profile upon T cell activation. Because the aptamer pull-down and MS analysis from nonactivated and activated T cells was performed simultaneously with CD3^+^ T cell lysates from the same donors (Fig. 2B, 3A), we could directly compare these two data sets. Interestingly, the interaction of AU-rich binding protein ZFP36L1 with all three cytokine 3’UTRs considerably increased after T cell activation (Fig. 3B). T cell activation also augmented the interaction of m6A-methylation reader YTH domain-containing family protein 1 (YTHDF1) to the *IFNG* 3’UTR, and of ribosomal RNA processing protein 1 homolog B (RRP1B) to the *TNF* 3’UTR. RBPs enriched for *IL2* 3′UTR included RRM-containing protein (RBM42), PC4 and SFRS1-interacting protein (PSIP1) and WARS (Fig. 3B). In contrast, ZC3H12D (Regnase-4) reduced its interaction to undetectable levels with all cytokine 3’UTRs upon T cell activation, as did uridylyl transferase ZCCHC11 and PABPN1 to the *IFNG* 3’UTR, ATP-dependent RNA helicase DDX39A and ZNHIT6 to the *TNF* 3’UTR, and for DDX46 and USO1 to the *IL2* 3’UTR (Fig. 3B). Again, several RBPs previously identified in an RBPome study (*26*) included HNRNPC, HuR, PUM2, PTBP2, CSDE1, and G3BP1.

Intriguingly, altered interactions with cytokine 3’UTRs correlated only in some cases with altered RBP expression levels upon T cell activation. MS analysis on total cell lysates of activated and nonactivated T cells (Dataset S5) revealed that the increased binding of ZFP36L1 to cytokine 3’UTRs coincided with increased protein expression in activated T cells, and reduced interaction of ZC3H12D with cytokine 3’UTRs coincided with decreased protein expression (Fig. 3C, SI Appendix, Fig. S2H). Most other RBPs, however, like WARS and YTHDF1 or DDX39A and DDX46 altered their binding to cytokine 3’UTRs without changing their protein expression levels (Fig. 3C, SI Appendix, Fig. S2H). We validated the protein expression levels for these RBPs in published data sets from reactivated memory T cells (*34*), and we observed increased expression levels only for ZFP36L1 to up to 24 h of T cell activation (Fig. 3D). In conclusion, T cell activation results in dynamic changes of the RBP binding landscape to cytokine 3’UTRs, which can only in part be explained by altered RBP protein expression.

### RBPs differentially modulate cytokine production in T cells

We next investigated if and how the identified RBPs modulated the cytokine production in human effector T cells. To this end, we deleted 5 strong interactors by CRISPR/Cas9 gene editing in α-CD3/α-CD28 activated human CD3^+^ T cells (Dataset S6). FXR2, HuR, ZC3HAV1, and ATXN2L interacted to cytokine 3’UTRs in both nonactivated and activated T cells, and ZFP36L1 increased its binding upon T cell activation. Of note, cell count and viability did not substantially differ from control cells that were nucleofected with non-targeting crRNA (SI Appendix, Fig. S3A). To mimic TCR-dependent activation, RBP-deleted T cells were activated for 4 h with α-CD3/α-CD28. Deleting FXR2 had no effect on cytokine expression when compared to donor-matched control CD8^+^ T cells (Fig. 4A, SI Appendix, Fig. S3B). In sharp contrast, reduced HuR expression diminished the cytokine production in CD8^+^ T cells (Fig. 4A, SI Appendix, Fig. S3B). Conversely, deleting ATXN2L, ZC3HAV1, or ZFP36L1 resulted in higher percentages of IFN-γ, TNF and IL-2 producing CD8^+^ T cells (Fig. 4A, B, SI Appendix, Fig. S3B). Native RNA-immunoprecipitation (RIP) with antibodies directed against ATXN2L, ZC3HAV1, HuR, and ZFP36L1 confirmed direct interaction of these RBPs with endogenous cytokine mRNA (Fig. 4C), indicating that these RBPs modulate cytokine production at least in part through direct interaction with the cytokine mRNA. The RIP essays also confirmed preferential binding of RBPs to specific cytokine mRNAs as shown in Fig. 2B and 3A, e.g. ATXN2L and ZFP36L1 to IL2 and HuR/ELAVL1 to TNF and IL-2 (cluster 7 and cluster 3, respectively; Fig. 2B).

**Fig. 4.**
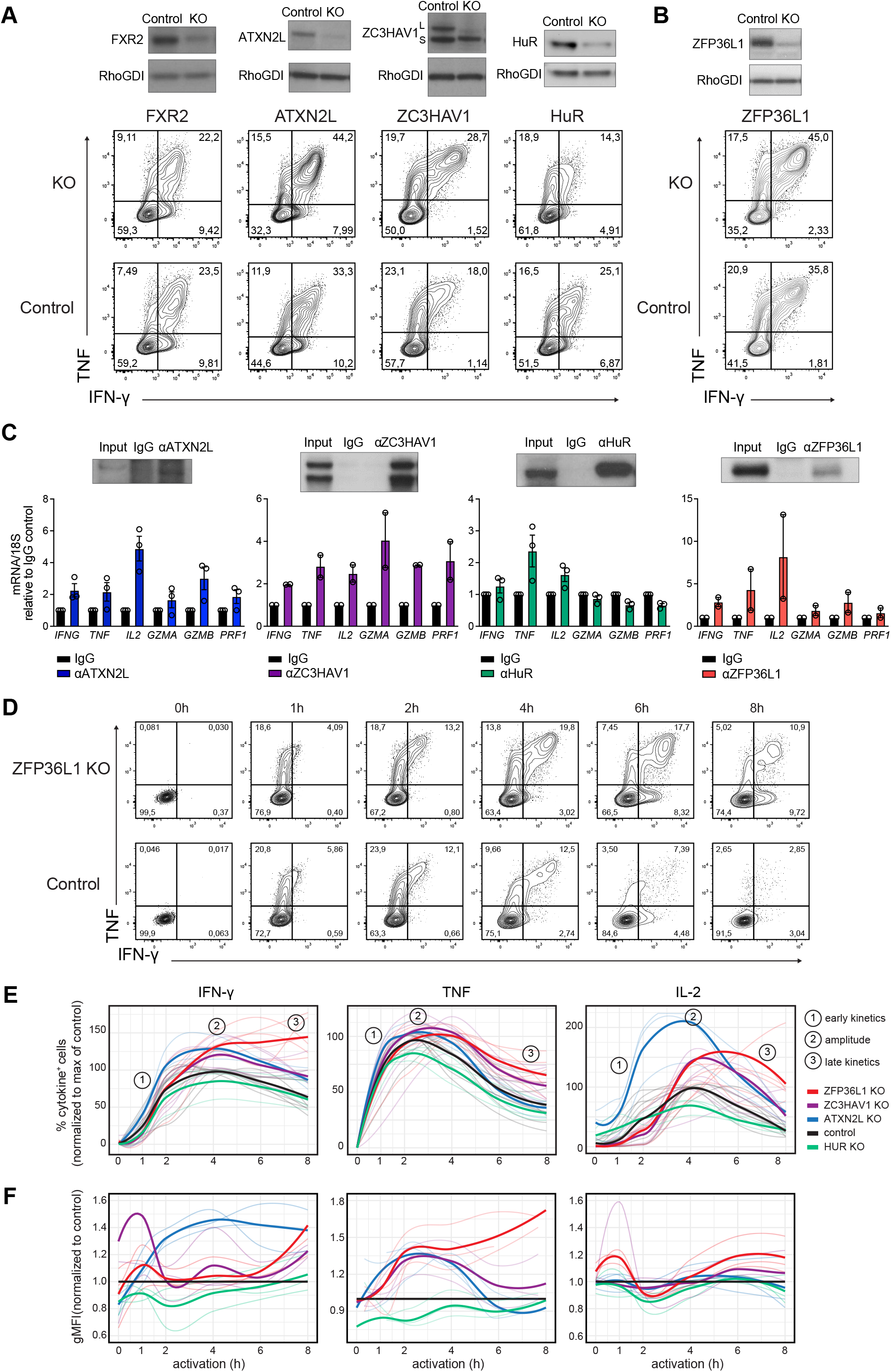
RBP-specific modulation of cytokine production kinetics in human T cells. (*A-B*) (top) KO efficiency of indicated RBPs in human T cells 7 days after CRISPR/Cas9 gene-editing. Non-targeting crRNA was used as control. L indicates long and S indicates short variant of ZC3HAV1. (bottom) IFN-γ and TNF protein expression in CD8^+^ T cells after 4 h of activation with α-CD3/α-CD28. Brefeldin A was added for the last 2 h. Data are representative of at least 2 independent experiments, each with 3 donors. (*C*) Native RNA immunoprecipitation (RIP) with antibodies specific for ATXN2L, ZC3HAV1, HuR, or ZFP36L1 from lysates of human T cells activated with PMA/I for 2 h (or 1 h for HuR). The respective IgG isotype was used as control. (top) Enrichment of indicated RBP upon RIP, as determined by immunoblot. (bottom) RBP interaction with endogenous mRNAs was determined by qRT-PCR. Compiled data of 3 donors for ATXN2L and HuR and 2 donors for ZC3HAV1 and ZFP36L1, from 2-3 independently performed experiments, presented as mean ± s.d. (*D*) IFN-γ and TNF production kinetics of nonactivated ZFP36L1 KO and control CD8^+^ T cells stimulated with α-CD3/α-CD28 for indicated time points. Brefeldin A was added for the last 2 h (or from the beginning for the 1 h time point). Representative dot plots as determined by intracellular cytokine staining. Data are representative of at least 6 donors from 2 independent experiments. (*E*) Cytokine protein production kinetics of human RBP KO CD8^+^ T cells stimulated with α-CD3/α-CD28 for indicated time points, compared to the peak of production (set at 100%) of the paired control treated T cell sample. Data are presented as mean (bold line) with individual donors (thin lines). (*F*) Cytokine protein production kinetics shown as gMFI of human RBP KO CD8^+^ T cells stimulated with α-CD3/α-CD28 for indicated time points compared to the control treated T cell sample. Data are presented as mean (bold line) with individual donors (thin lines).

### RBPs display different kinetics in modulating cytokine production

We and others previously showed that IFN-γ, TNF, and IL-2 follow individual production kinetics in CD8^+^ T cells (*12, 35, 36*). To investigate how RBP depletion influences the cytokine production kinetics, we reactivated CD3^+^ T cells with α-CD3/α-CD28 for 1 h to up to 8 h, and measured cytokine production, by adding brefeldin A for a maximum of 2 h (*12*). This snapshot analysis allowed to define the activity kinetics of RBPs and their mode of action, as exemplified by ZFP36L1-deficient T cells: ZFP36L1 KO CD8^+^ T cells show similar cytokine production profiles in the first two hours, but increase their production profile in particular at later time points (Fig. 4D).

When we followed the cytokine production kinetics of RBP KO T cells, we observed a time-dependent and RBP-specific contribution to the cytokine production kinetics. HuR-deficient CD8^+^ T cells showed reduced cytokine production upon 1-4 h of activation (Fig. 4E, SI Appendix, Fig. S3D-E), a feature that was only observed in CD8^+^ T cells, but not in CD4^+^ T cells (Fig. 4E and SI Appendix, Fig. S4). Conversely, ATXN2L depletion augmented the cytokine production in CD8^+^ and CD4^+^ T cells early on, and did so most effectively for IL-2, where the peak of response was also earlier than that of control cells (Fig. 4E, SI Appendix, Fig. S3D-E and Fig. S4). Differences in IFN-γ production was most prominent in the per cell basis of production, as defined by the gMFI (Fig. 4F). Yet again, the cytokine production kinetics at later time points after the peak of the response followed the same slope as control T cells (Fig. 4E, SI Appendix, Fig. S3D-E and Fig. S4). ZC3HAV1 deletion primarily increased the cytokine production in CD4^+^ and CD8^+^ T cells at the peak of the response, exceeding that of control T cells and from 4 h of activation onwards, but with no alterations in the kinetics of cytokine production (Fig. 4E, SI Appendix, Fig. S3D-E).

ZFP36L1 deletion showed identical response rates in CD8^+^ T cells as ZC3HAV1 in the first 4 h of activation. However, ZFP36L1 deletion was the only RBP where the slope of reduced cytokine production at later time points diverged from the other RBP KO cells. In fact, high cytokine production was maintained to up to 8 h post T cell activation (Fig. 4D-E, SI Appendix, Fig. S3D-E). Interestingly, ZFP36L1 KO CD8^+^ T cells showed the largest differences for IFN-γ and IL-2 production in percentages of cytokine-producing T cells, whereas the TNF production primarily differed in the magnitude of production, as determined by the gMFI (Fig. 4D-F). The effects of ZFP36L1 depletion were also observed in CD4^+^ T cells, but less so (SI Appendix, Fig. S4). In conclusion, the four tested RBPs regulate cytokine production in human T cells in an RBP-specific manner, displaying different effects in regulating the onset, magnitude and duration of cytokine production (SI Appendix, Fig. S3F).

### ZFP36L1 destabilizes cytokine mRNA in T cells

To investigate how RBPs modulate cytokine production, we measured their effect on mRNA stability. For ZFP36L1, ATXN2L and ZC3HAV1 KO CD3^+^ T cells, we activated the cells for 3h with α-CD3/α-CD28 and treated them with Actinomycin D (ActD) to block *de novo* transcription. Due to its different activity kinetics, we started the ActD treatment for HuR KO cells earlier, i.e. at 1 h post T cell activation. Intriguingly, whereas deletion of ATXN2L, ZC3HAV1 or HuR had no effect on RNA stability or overall mRNA levels (Fig. 5A, SI Appendix, Fig. S5A), ZFP36L1 deletion substantially increased the mRNA half-life of all three cytokine mRNAs: for *IFNG* and *IL2* from 50-60 min in control-treated T cells to >120 min, and for *TNF* from 20-30 min to 40-50 min in ZFP36L1 KO T cells (Fig. 5A). The increased mRNA stability also resulted in higher overall cytokine mRNA expression levels in activated ZFP36L1 KO T cells compared to control T cells (Fig. 5B). A co-immunoprecipitation assay with α-ZFP36L1 antibodies from activated human CD3^+^ T cell lysates followed by MS analysis revealed that ZFP36L1 interacts with many components of the mRNA degradation machinery in T cells, including CNOT3, CNOT4, CNOT6L, CNOT7, CNOT10, CNOT11 from the CCR4-NOT complex, and EXOSC1, EXOSC2, EXOSC5, EXOSC6, EXOSC7, EXOSC8, EXOSC9 from the RNA 3’->5’ exosome complex (Fig. 5C, SI Appendix, Fig. S5B). Thus, ZFP36L1 dampens cytokine production in T cells by destabilizing its target mRNA.

**Fig. 5.**
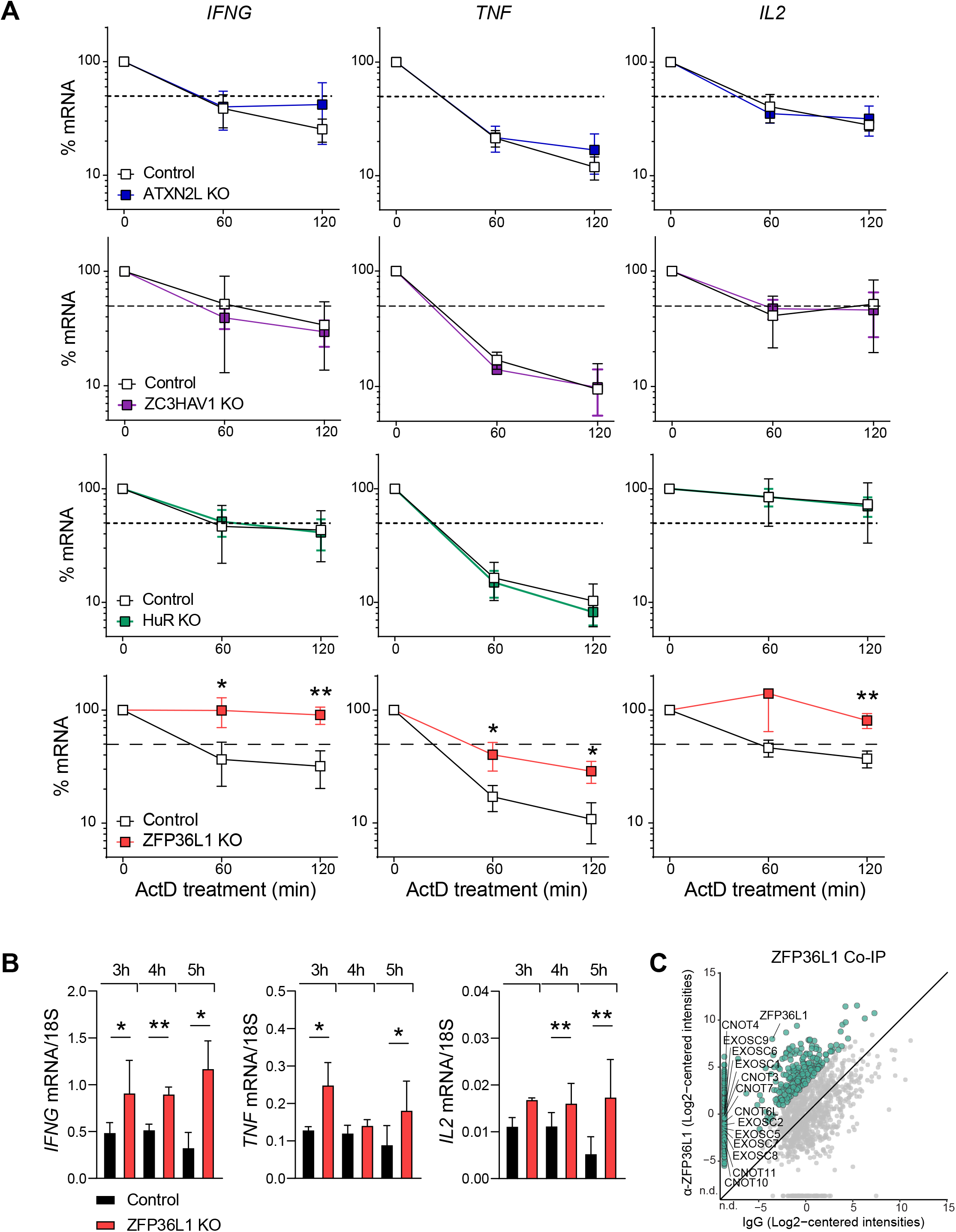
ZFP36L1 destabilizes IFN-γ, TNF and IL-2 mRNA in human T cells. (*A*) *IFNG, TNF* and *IL2* mRNA levels of ATXN2L, HuR, ZC3HAV1, and ZFP36L1 KO T cells and paired control-treated T cells that were reactivated with α-CD3/α-CD28 for 3h (HuR for 1 h), and that were then treated with Actinomycin D (ActD) for indicated time points. Data are presented as mean ± s.d. of 3 donors and are representative of 2 independently performed experiments. Unpaired t test (*P <0.05; **P<0.01). (*B*) Cytokine mRNA levels of ZFP36L1 KO and control T cells that were reactivated for indicated time points with α-CD3/α-CD28. Data are presented as mean ± s.d. of 3 donors and are representative of 2 independently performed experiments. Ratio paired t test (*P <0.05; **P<0.01). (*C*) Co-immunoprecipitation with α-ZFP36L1 antibodies or IgG control from cytosolic lysates of 2 h PMA/I-activated human T cells, followed by MS analysis. Data represent protein raw log2 median-centered intensities from ZFP36L1 immunoprecipitation versus control. Expression levels of biological replicates were averaged (n=3 donors). Green dots depict putative interactors of ZFP36L1 that were identified with cut-off LFC>6 compared to control. Gray dots depict all proteins identified in one experiment. n.d.= not detected.

### RBPs define the cytokine production of T cells in response to target cells

We next defined the effect of RBP depletion in T cells when responding to tumor target cells. We first retrovirally transduced T cells with a MART1-specific TCR recognizing the HLA-A*0201-restricted MART1_26-35_ epitope (*37*) and then deleted RBPs that block cytokine production by CRISPR/Cas9 gene editing (SI Appendix, Fig. S6A). The RBP-deficient, or control treated MART1 TCR-expressing T cells were then exposed for different time points to a MART1^hi^ HLA-A*0201^+^ melanoma cell line (MART1^+^), or to a MART1^lo^ HLA-A*0201^-^ melanoma control cell line (MART1^-^) (SI Appendix, Fig. S6B). At early time points (2 and 4 h) after co-culture, ATXN2L KO CD8^+^ T cells showed higher IL-2 and TNF production, and ZC3HAV1 KO T cells displayed a slight, but significant increase in TNF and IFN-γ producing CD8^+^ T cells at 4 h (SI Appendix, Fig. S6C). At later time points, ZFP36L1 KO CD8^+^ T cells were superior cytokine producers (Fig. 6A-C). As previously reported (*35, 36*), all T cells showed reduced cytokine production at 24 h of co-culture, in particular of IL-2 and TNF. At this time point, ZFP36L1 KO CD8^+^ T cells most robustly maintained the cytokine production (Fig. 6B, C). Furthermore, ZFP36L1 KO CD8^+^ T cells maintained their capacity to co-produce several cytokines (Fig. 6B), a feature that is indicative for the most-potent anti-viral and anti-tumoral CD8^+^ and CD4^+^ T cells in humans (*3, 4*). This higher cytokine production -albeit to a lesser extent -was also observed for ATXN2L KO CD8^+^ T cells, which primarily maintained their capacity to produce IFN-γ (Fig. 6B). Of note, RBP deletion did not substantially modulate the expression of surface molecules, such as CD3, CD69, CD137 or PD-1 (SI Appendix, Fig. S6D). In conclusion, depleting individual RBPs can boost and prolong cytokine production of T cells in response to target cells.

**Fig. 6.**
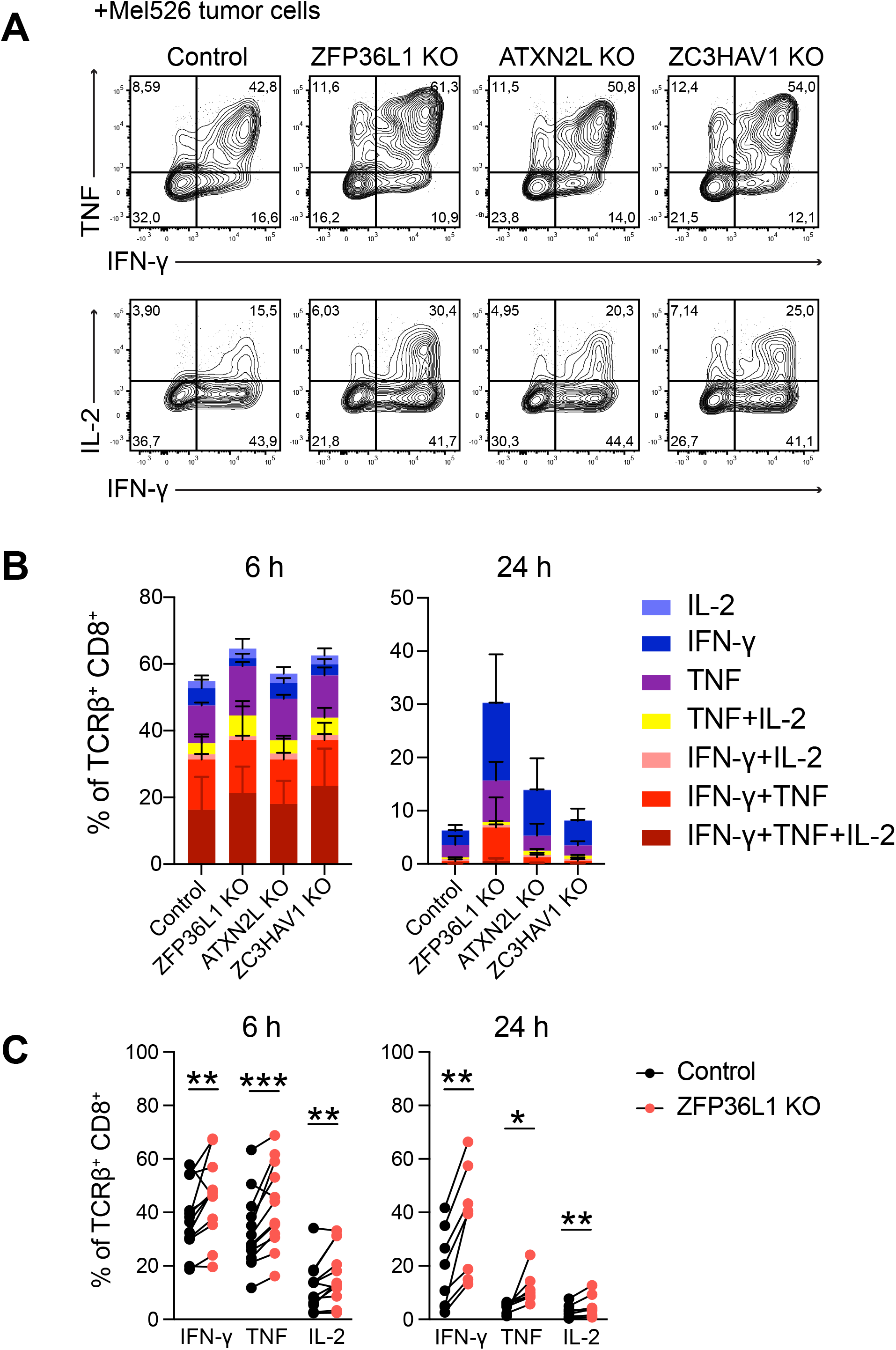
RBPs regulate cytokine expression in tumor-specific human T cells. (*A*) MART1 TCR-engineered T cells were CRISPR/Cas9 gene-edited for indicated RBP, or treated with non-targeting crRNA as control. Representative dot plots of IFN-γ and TNF production in CD8^+^ T cells after 6 h of co-culture with MART1^+^ tumor cells. Brefeldin A was added for the last 2 h. Data are representative of at least 2 independent experiments, each with 3-4 donors. (*B*) Graphs depict the cytokine profile analysis of MART1-specific CD8^+^ T cells. Data are presented as mean ± s.d. of 3 donors and are representative of at least 2 independent experiments. (*C*) Compiled data of 7 donors after 6 and 24 h of stimulation with MART1^+^ tumor cells from 2 independently performed experiments. Data are presented as mean ± s.d. Ratio paired t test (*P <0.05; **P<0.01; ***P<0.001).

### Superior anti-tumoral responses by ZFP36L1-deficient T cells *in vivo*

Gradual loss of effector function, and in particular of IFN-γ and TNF is a major hurdle of effective T cell responses to tumors (*38, 39*). Furthermore, we previously showed that post-transcriptional mechanisms block IFN-γ production in murine TILs, but which RBPs mediate this regulation remains unknown (*16*). Thus, to study whether RBP deletion rescues cytokine production of TILs *in vivo*, we turned to mouse models. We focused our attention on ZFP36L1 and ATXN2L, which showed the most potent effects in human T cells (Fig. 6). We generated murine Zfp36l1- and Atxn2l-deficient OT-I T cells by CRISPR/Cas9 gene-editing. Whereas deletion of Atxn2l in OT-I T cells failed to recapitulate its role in regulating cytokine production in murine T cells (SI Appendix, Fig. S7A, B), Zfp36l1 KO OT-I T cells showed superior IFN-γ production to control cells upon 4 h of activation with OVA_257-264_ peptide (SI Appendix, Fig. S7A, B). Thus, the regulation of cytokine production by ZFP36L1 is conserved.

To determine the role of ZFP36L1 *in vivo*, we injected 1×10^6^ OT-I Zfp36l1 KO, or control OT-1 T cells into tumor-bearing mice that had received OVA-expressing B16 melanoma cells (B16-OVA; Fig. 7A) 7 days earlier. 14 days post T cell transfer, we first analyzed the cytokine production of T cells in the spleen. The percentage of IFN-γ and IL-2 production in Zfp36l1 KO OT-I T cells was slightly, but not significantly increased compared to control OT-I T cells (SI Appendix, Fig. S7C). We then turned our attention to tumor-infiltrating T cells. The absolute number of Zfp36l1 KO and control OT-I TILs did not substantially differ (Fig. 7B), indicating that the Zfp36l1-deficiency had no overt effects on survival and/or proliferation. Also, the PD-1 expression was consistently high in control and Zfp36l1 KO OT-I T cells (SI Appendix, Fig. S7D). Nonetheless, Zfp36l1 KO OT-I TILs showed slightly higher percentages of Tcf1^-^Tox^+^ terminally differentiated dysfunctional T cells and reduced percentages and significantly lower expression levels of Tcf1^+^ progenitor dysfunctional T cells (Fig. 7C). Concomitant with this shift in transcription factor expression, Zfp36l1 KO OT-I TILs displayed lower Slamf6 expression and increased Tim-3 expression (Fig. 7D), demonstrating that Zfp36l1 deletion could not rescue the dysfunctional state of T cells.

**Fig. 7.**
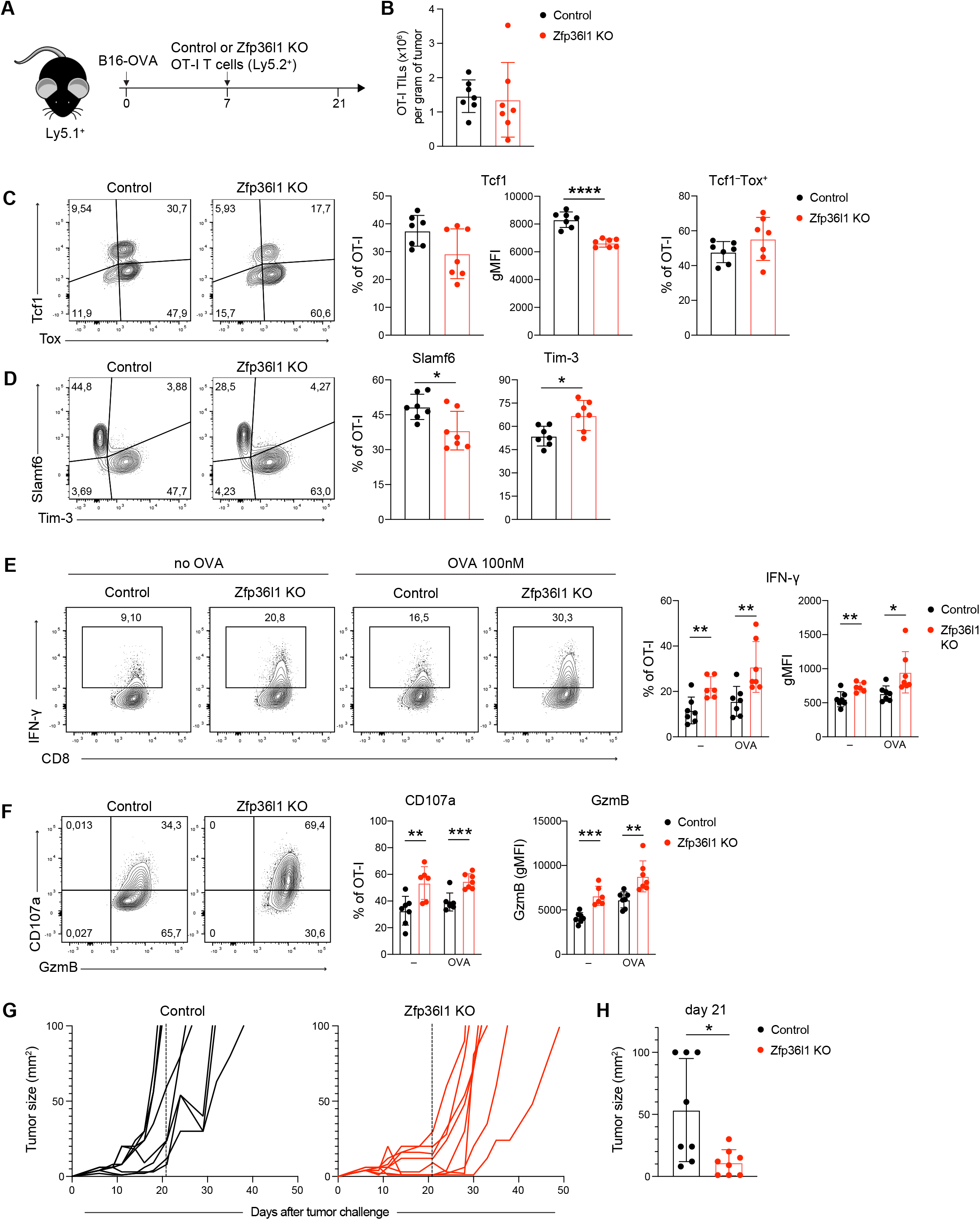
ZFP36L1 dampens cytokine production in tumor-specific T cells *in vivo*. (*A*) Ly5.1^+^ C57BL/6J mice were engrafted with B16-OVA tumors. 7 days later mice received 1×10^6^ control or Zfp36l1 KO OT-I Ly5.2^+^ T cells. 14 days after T cell transfer, tumors were excised and tumor-infiltrating T cells (TILs) were analyzed. (*B*) Absolute numbers of control and Zfp36l1 KO OT-I TILs per gram of tumor. Unpaired t test. (*C-D*) (left) Representative protein expression of transcription factors (Tcf1, Tox) and surface markers (Slamf6, Tim-3) by control or Zfp36l1 KO OT-I TILs. (right) Percentages or gMFI of transcription factors and surface markers-expressing OT-I TILs. Data are presented as mean ± s.d. with n=7, and are representative of 2 independent experiments. Unpaired t test (*P <0.05; ****P <0.0001). (*E*) (left) Representative IFN-γ protein expression by control or Zfp36l1 KO OT-I TILs ex vivo with or without reactivation with 100nM OVA_257–264_ peptide for 4 h, brefeldin A and monensin added for the last 2 h. (right) Percentage or gMFI of IFN-γ producing OT-I CD8^+^ T cells. Data are presented as mean ± s.d. with n=7, and are representative of 2 independent experiments. Unpaired t test (*P <0.05; **P<0.01). (*F*) (left) Representative CD107a and Granzyme B expression by control and Zfp36l1 KO OT-I TILs ex vivo in the presence of brefeldin A and monensin for 2 h. (right) Zfp36l1 KO and control OT-I T cells reactivated with 100nM OVA_257–264_ peptide for 4 h, or left untreated, in the presence of brefeldin A and monensin for the last 2 h. Data are presented as mean ± s.d. with n=7 and are representative of 2 independent experiments. Unpaired t test (**P<0.01; ***P<0.001). (*G*) Tumor size of Ly5.1^+^Ly5.2^+^ B16-OVA tumor-bearing mice treated with 0.65×10^6^ Zfp36l1 KO or control OT-I Ly5.2^+^ T cells at day 7 after tumor injection. Dashed line represents time point of 14 days after the T cell transfer. n = 8 mice/group. (*H*) Tumor size of tumor-bearing mice at day 21 (14 days after the T cell transfer). n = 8 mice/group. Unpaired t test (*P<0.05).

Despite their phenotypical appearance of dysfunctional cells, Zfp36l1 KO TILs were superior cytokine producers. Control OT-I TILs failed to produce detectable levels of TNF and IL-2, and very limited production of IFN-γ was measured when tumor digests were incubated *ex vivo* for 2 h with brefeldin A and monensin without the addition of exogenous peptide (Fig. 7E, and SI Appendix, Fig. S7E) (*16*). Restimulation for 4 h with OVA_257-264_ peptide only marginally increased the cytokine expression in control OT-I TILs, in particular when compared to reactivated splenic OT-I T cells (Fig. 7E and SI Appendix, Fig. S7C, E). In contrast, Zfp36l1 KO OT-I TILs not only produced significantly more IFN-γ when cultured with or without exogenous antigen (Fig. 7E), they also showed increased production of TNF and IL-2 (SI Appendix, Fig. S7E). This was true both for the percentage of cytokine-producing TILs and for the cytokine production per cell (Fig. 7E and SI Appendix, Fig. S7E). Intriguingly, also other effector molecules such as the degranulation marker CD107a and the cytotoxic molecule Granzyme B were increased in Zfp36l1 KO TILs compared to control OT-I TILs (Fig. 7F). This feature appeared to be unique to the tumor environment, because no difference in CD107a and Granzyme B expression was observed in spleen-derived OT-I T cells (SI Appendix, Fig. S7F).

Because Zfp36l1 KO TILs displayed increased cytokine production and granzyme B expression, we hypothesized that Zfp36l1-deficiency could also influence the tumor outgrowth. We therefore measured the tumor growth over time in B16-OVA tumor-bearing mice that had received 0.65×10^6^ Zfp36l1 KO or control OT-I T cells. Remarkably, even these low numbers of transferred Zfp36l1 KO OT-I T cells could significantly delay the tumor outgrowth of this aggressive tumor (Fig. 7G). In fact, at 14 days post T cell transfer, i.e. the time point when we measured superior cytokine production and degranulation in Zfp36l1 KO TILs *ex vivo* (Fig. 7G, H), 3/8 of the control mice and none of the Zfp36l1 KO mice reached the human endpoint of 1000mm^3^ of tumor size (Fig. 7H). In conclusion, Zfp36l1 gene-editing in T cells results in superior cytokine producers in the tumor environment, which enhances the therapeutic potency of T cell therapy.

## Discussion

Our study uncovers the breath of RBP interactions with cytokine 3’UTRs in human T cells. These RNA-RBP interactions are subject to alterations upon T cell activation. For ZFP36L1 and ZC3H12D, these alterations in RNA binding coincides with altered RBP protein expression. However, most RBPs maintain their expression profile upon T cell activation. It is therefore reasonable to consider that post-translational modifications contribute to these altered binding activities. For instance, stress-induced phosphorylation of HuR reduced its interaction with target mRNA in HeLa cells (*40, 41*). Future studies that allow combined interrogation of the interactome and the phosphoproteome may shed light on such alterations (*42*). Alternatively, RBPs can also compete for binding, as shown for Arid5a and ZC3H12A on the *STAT3* 3’UTR, and for HuR and ZFP36 (Tristetraprolin, TTP) on *TNF* and *GM-CSF* mRNA (*20, 43, 44*). Lastly, other RNA regulatory molecules such as miRNAs can compete with RBPs (*45*). Understanding these mechanisms that define RBP activity will further help decipher the role of RBPs in modulating T cell responses.

Overall, we identified between 60-90 RBPs interacting with each cytokine 3’UTR, a number that corresponds well with reports from other RNA-centric RBP identification approaches with different RNA target (*46*). It requires further investigation whether these numbers stem from ample RBP binding to each RNA molecule, or from heterogeneous RBP-RNA interactions within one T cell, which may result from different subcellular RNA (and RBP) localizations. It could also stem from heterogenous expression of RBPs within the T cell population used for bait, due to the non-synchronous state found even in cultured T cells. Importantly, while the identification of some RBPs interacting with unmodified 3’UTRs such as PABPs, ERI1 or YTHDF1 may sound counterintuitive, it is very well representing the biological activity of these proteins. Not only are putative binding motifs for these RBPs found in cytokine 3’UTRs (Fig. 1D), but also does YTHDF1 interact with non-methylated RNA, albeit with a lower affinity (*47*). Because the pulldown was performed on whole-cell T cell lysates, it is yet to be defined at which subcellular localization the mRNA-RBP interactions occurred, or whether nuclear RBPs translocated to the cytosol upon T cell activation. In sum, our study highlights the potential to screen for RNA-RBP interactions from full length 3’UTRs, forming the basis to further decipher the functional consequences of these interactions. Of note, due to the stringent cut-off of LFC>4 we used, the provided RBP list may not be complete. Nevertheless, it also holds less contaminants. RBP candidates with lower confidence are available in Datasets S1, S3 and can be further investigated. As for all such screens alike, all identified RBPs require further validation irrespective of the confidence threshold employed, through e.g. RIPs to measure interactions with endogenous RNAs, and through defining their functional activity and mode of action through e.g. genetic modifications.

Validation studies of RBPs identified in the pull-down assay revealed that four out of five tested RBPs modulate cytokine production in human T cells. Intriguingly, we observed RBP-specific effects on cytokine production, including their mode of action and their activity kinetics. HuR promotes cytokine production in human T cells at the onset (1-2 h) of T cell activation. This early boost in cytokine production may in particular be relevant for antigen-experienced T cells, which contain pre-formed mRNA, and for which immediate cytokine production is licensed by releasing the ready-to deploy mRNA from ZFP36L2 (*12, 22*). We here found that this early cytokine production is further facilitated by HuR. How HuR regulates cytokine production is yet to be defined. Because we found no effects on mRNA stability, we postulate that HuR influences translation directly, or indirectly by rendering mRNA accessible to the translation machinery through altering its subcellular localization, or through competing with other RBPs.

ZFP36L1, ATXN2L, and ZC3HAV1 block cytokine production most effectively at 4-8 h post activation, with ZFP36L1 (and ATXN2L) also at 24 h of T cell activation. ATXN2L and ZC3HAV1 do not affect RNA stability, and thus employ other post-transcriptional events to dampen cytokine production. In epithelial cell lines, ATXN2L regulates stress granules and processing bodies (*48*). ZC3HAV1 destabilizes *IFNB and IFNL3* mRNAs in hepatocytes and human cytomegalovirus RNA in fibroblasts and inhibits programmed ribosomal frameshifting of the SARS-CoV-2 virus (*49–51*). The mechanisms that ATXN2L and ZC3HAV1 employ in T cells to suppress cytokine production, however, may differ and await further elucidation. ZFP36L1 destabilizes all 3 key cytokines produced by effector CD8^+^ T cells, and it does so at least in part through target mRNA degradation. ZFP36L1 was shown to interact with CNOT7, a component of the CCR4-NOT mRNA degradation complex in 293T cells (*52*), and CNOT7 degraded ZFP36 targets in HeLa cells (*53, 54*). We here extended these findings of ZFP36L1 interactions to many other members of the CCR4-NOT complex and to several exonucleases. In sum, all three negative regulators of cytokine production, i.e. ZFP36L1, ATXN2L, and ZC3HAV1 highlight their critical contribution in preventing excessive production of cytokines in activated T cells. Having multiple RBP regulators with different modes of action and different activity kinetics also further highlights the necessity of fine-tuning the production of cytokines for appropriate T cell function.

Combined deletion of ZFP36 family members was previously shown in mouse models to modulate T cell responses to infection (*55*). Here, we report that single deletion of ZFP36L1 in human T cells augments cytokine production, and it delays the shutdown of cytokine production when T cells are exposed to tumor cells. Importantly, T cells lose their capacity to produce cytokines within the tumor environment, a process we previously showed to rely on post-transcriptional events (*16, 17*). In this study, we found that deleting ZFP36L1 boosts the effector function of TILs *in vivo*, which resulted in superior anti-tumor responses. ZFP36L1 deletion does not only improve the production of IFN-γ, but also that of TNF and IL-2, and that of cytotoxic mediators, Granzyme B and CD107a, indicating that ZFP36L1 instructs several T cell effector programs. Therefore, ZFP36L1 represents an attractive target to boost anti-tumoral T cell effector responses.

In sum, we here provide the first catalogue of dynamic RBP interactions to cytokine mRNAs in human T cells. Our results reveal that this map can serve as valuable resource for studying the role of RBPs in regulating T cell responses to identify RBPs as potential targets for therapeutic purposes. Considering the association of RBPs with human genetic disorders (*56*), our work should pave the way to support further studies on the role of RBPs in other immune-related disease settings.

## Materials and methods

### Cloning and preparation of *in vitro* transcribed S1m aptamers

Full length 3′UTRs were amplified from human, or from C57BL/6J mouse-derived genomic DNA and cloned into BamHI and NotI sites of pRETRO-SUPER GFP (*57*) downstream of GFP (Dataset S6). 4xS1m RNA aptamers containing the full length 3’UTR of human *IFNG, TNF* and *IL2* were cloned into the pSP73-4xS1m vector (*31*) and *in vitro* transcribed as previously described (*22*). RNA quality and quantity was determined by RNAnano Chip assay (Agilent).

### Mice

C57BL/6J/Ly5.1, C57BL/6J/Ly5.1/Ly5.2 and C57BL/6J.OTI T cell receptor (TCR) transgenic mice (OT-I) mice were bred in-house at the Netherlands Cancer Institute (NKI). Experiments were performed with 6-12 week-old mice in accordance with institutional and national guidelines and approved by the Experimental Animal Committee at the NKI.

### T cell activation and culture

Human T cells from anonymized healthy donors were used in accordance with the Declaration of Helsinki (Seventh Revision, 2013) after written informed consent (Sanquin). Peripheral blood mononuclear cells (PBMCs) were isolated through Lymphoprep density gradient separation (Stemcell Technologies). Cells were used after cryopreservation. Human T cells were activated for 48 h as previously described (*58*), harvested and further cultured in standing T25/75 tissue culture flasks (Thermo Scientific) at a density of 0.8×10^6^/mL in IMDM supplemented with 5% heat-inactivated human serum (Sanquin), 5% FBS and 100 IU/mL recombinant human (rh) IL-2 (Proleukin, Clinigen). Medium was refreshed every 3 days. Upon nucleofection, T cells were cultured in T-cell mixed media (Miltenyi) supplemented with 5% heat-inactivated human serum, 5% FBS, 100 U/mL Penicillin, 100 μg/mL streptomycin, 2 mM L-glutamine, 100 IU/mL rhIL-2 and 10 ng/mL rhIL-15 (Peprotech). OT-I T cells were purified from spleens with the CD8a^+^ T Cell Isolation Kit (Miltenyi Biotec), activated and cultured as previously described (*11*).

### Retroviral transduction

Transduction of T cells was performed with Retronectin (Takara) as previously described (*22, 58*).

### 4xS1m RNA aptamer-protein pull-down

α-CD3/α-CD28 activated human CD3^+^ T cells were rested for 5 days. T cells activated with PMA/Ionomycin for 2 h or that were left untreated were washed twice with ice-cold PBS. Cell pellet was snap frozen in liquid nitrogen, homogenized using 5-mm steel beads and a tissue lyser (Qiagen TissueLyser II) 6x at 25 Hz for 15s. The homogenate was then solubilized and precleared twice as previously described (*22*). Cell lysates were incubated with RNA-aptamer-coupled beads for 3.5 h at 4°C under rotation in the presence of 60U RNasin (Ambion). For each pull-down, 30 μg of *in vitro* transcribed RNA, coupled to Streptavidin Sepharose High Performance beads (GE Healthcare), and 5-10 mg cell lysate protein was used. RNA-bound proteins were eluted by adding 1 μg RNaseA (Thermo Scientific) in 100 μl 100 mM Tris-HCl, pH 7.5 (Gibco-Invitrogen). Proteins were reduced, alkylated and digested into peptides using trypsin (Promega). Peptides were desalted and concentrated using Empore-C18 StageTips and eluted with 0.5% (v/v) acetic acid, 80% (v/v) acetonitrile. Sample volume was reduced by SpeedVac and supplemented with 2% acetonitrile and 0.1% TFA.

### Co-immunoprecipitation

Cytoplasmic lysates of 100×10^6^ PMA/Ionomycin-activated human CD3^+^ T cells were prepared using lysis buffer (140 mM NaCl, 5 mM MgCl_2_, 20 mM Tris/HCl pH7.6, 1% Digitonin) freshly supplemented with 1% of protease inhibitor cocktail (Sigma). Protein A Dynabeads (Invitrogen) were prepared according to the manufacturer’s protocol. The lysate was immunoprecipitated for 4 h at 4°C with 10 μg polyclonal rabbit α-ZFP36L1 (ABN192, Sigma-Aldrich) or with isotype control (12-370, Sigma-Aldrich). Beads were washed twice with wash buffer (150 mM NaCl, 10 mM Tris/HCl pH7.6, 2 mM EDTA, protease/phosphatase inhibitor cocktail) and twice with 10 mM Tris/HCl pH7.6. Immunoprecipitated proteins were reduced and on-bead alkylated. Proteins were detached with 250 ng trypsin for 2 h at 20°C. Beads were removed and proteins were further digested into peptides with 350 ng trypsin for 16 h. Peptides were prepared for MS analysis, as described above.

### Mass spectrometry analysis

Mass spectrometry data acquisition was performed as described in *SI*. Raw mass spectrometry files were processed with the MaxQuant computational platform, version 1.6.2.10 (*59*). Proteins and peptides were identified using the Andromeda search engine by querying the human Uniprot database (downloaded February 2017 and February 2019, 89,796 entries). Standard settings with the additional options match between runs, and unique peptides for quantification were selected. The generated ‘proteingroups.txt’ data were imported in R and processed with the Differential Enrichment analysis of Proteomics data (DEP) R package (*60*). Identified peptides were filtered for potential contaminants, only identified by site and reverse hits.

### Genetic modification of T cells with Cas9 RNPs

crRNAs were designed in Benchling (https://benchling.com; Dataset S6). Cas9 RNP production and T cell nucleofection was performed as previously described (*17*). For nucleofection, human CD3^+^ T cells were activated for 72 h with α-CD3/α-CD28, and mouse OT-I T cells for 20h with MEC.B7.SigOVA cells (*61*) and rested for 24 h in medium with IL-7. Cells were electroporated in 16-well strips in a 4D Nucleofector X unit (Lonza) with program EH100 for human T cells, and with program CM137 for mouse T cells. Nucleofection efficiency was determined on day 2 after electroporation by measuring ATTO550 expression using FACSymphony (BD Biosciences). Knockout efficiency was determined on day 7-10 after electroporation by Western blot.

### Quantitative PCR analysis

Total RNA was extracted using Trizol (Invitrogen). cDNA was synthesized with SuperScript III (Invitrogen), RT-PCR was performed using SYBR Green on a StepOne Plus (Applied Biosystems). Ct values were normalized to 18S levels (*36*). For mRNA half-life measurements, T cells were activated in triplicate for indicated timepoint with α-CD3/α-CD28, and then treated with 5 μg/ml actinomycin D (ActD) (Sigma-Aldrich).

### RNA immunoprecipitation and immunoblotting

Cytoplasmic lysates of 300×10^6^ PMA/Ionomycin activated human CD3^+^ T cells were prepared as previously described (*22*). Protein A or protein G Dynabeads (Invitrogen) were prepared according to manufacturer’s protocol. The lysate was immunoprecipitated for 4 h at 4°C with 10 μg polyclonal rabbit α-ZFP36L1 (ABN192, Sigma-Aldrich), α-ZC3HAV1 (GTX120134, GeneTex), α-ATXN2L (ab99304, Abcam) or a polyclonal rabbit IgG isotype control (12-370, Sigma-Aldrich) and a mouse monoclonal α-HuR (3A2, Santa Cruz Biotechnology) or a mouse IgG1 kappa isotype control (P3.6.2.8.1, eBioscience). RNA was extracted directly from beads by using Trizol, and mRNA expression was measured by RT-PCR as described above. Specificity of the RNA-IP assay was tested by immunoblotting using α-ZFP36L1 (ab42473, Abcam), α-ZC3HAV1 (PA5-31650, Invitrogen), α-ATXN2L (ab99304, Abcam) or α-HuR (3A2, Santa Cruz Biotechnology), followed by mouse monoclonal anti-rabbit IgG light chain-HRP (ab99697, Abcam) or goat anti-mouse-HRP (1031-05, Southern Biotech).

### Functional assays

For *in vitro* assays, human CD3^+^ T cells were stimulated with 10ng/ml PMA and 1μM Ionomycin (Sigma-Aldrich) or with 1 μg/mL pre-coated α-CD3 and 1 μg/mL soluble α-CD28. MART1 TCR-transduced CD3^+^ T cells were co-cultured with HLA-A*0201^+^ MART1^hi^ Mel526 (MART1^+^) or HLA-A*0201-MART1^lo^ Mel888 (MART1^-^) melanoma cells (*58, 62*), in a 1:1 effector to target (E:T) ratio. 1 μg/mL brefeldin A (BD Biosciences) was added as indicated. Non-activated T cells were used as control.

### B16 melanoma tumor model

C57BL/6J/Ly5.1 or C57BL/6J/Ly5.1/Ly5.2 mice were injected subcutaneously with 1×10^6^ B16-OVA cells (*63*). On day 7 when tumors reached ∼4-8mm^2^, 0.65-1×10^6^ control or Zfp36l1 KO CD8^+^ OT-I Ly5.2 T cells were injected intravenously. Prior to T cell transfer, dead cells were removed with Lympholyte M gradient (Cedarlane). Tumor infiltrates were analyzed 14 days after T cell transfer. Excised tumors were cut into small pieces and digested at 37°C for 30 min with 100 μg/ml DNase I (Roche) and 200 U/ml Collagenase (Worthington). Cells were counted and incubated for 2 h with brefeldin A, or for 4 h with 100 nM OVA_257–264_ peptide and brefeldin A for the last 2 h of activation. For studying tumor outgrowth, mice were sacrificed when tumor reached ∼100 mm^2^.

### Flow cytometry and intracellular cytokine staining

T cells were washed with FACS buffer (PBS, containing 1% FBS and 2 mM EDTA) and labeled for 20min at 4°C with α-CD4 (SK3 and RPA-T4), α-CD8 (SK1), α-CD69 (FN50; all BD Biosciences), α-mouse TCR beta (H57-597), α-IFN-γ (4S.B3, both eBioscience), α-CD3 (OKT3), α-CD279 (EH12.2H7), α-CD137 (4B4-1), α-IL-2 (MQ1-17H12, all Biolegend), and α-TNF (MAb11, Biolegend and BD Biosciences). Mouse T cells were labelled with α-CD8 (53-6.7), α-PD-1 (J43), α-GzmB (GB11) (all BD Biosciences), α-CD45.1 (A20), α-CD45.2 (104), α-IFN-γ (XMG1.2) and α-TNF (MP6-XT22), α-IL-2 (JES6-5H4), α-CD107a (1D4B) (all eBioscience). Dead cells were excluded with Near-IR (Life Technologies). For intracellular cytokine staining, cells were cultured with 1 μg/mL brefeldin A for indicated timepoints, fixed and permeabilized with Cytofix/Cytoperm kit (BD Biosciences) prior to acquisition using FACSymphony. Data were analyzed with FlowJo (BD Biosciences, version 10).

## Statistical analysis

Results are shown as mean ± s.d. Statistical analysis was performed with GraphPad Prism 8 with two-tailed ratio paired or unpaired Student t test when comparing two groups, or with one-way ANOVA test with Dunnett correction when comparing more than two groups. p values <0.05 were considered statistically significant.

## Acknowledgements

We thank the animal caretakers of the NKI and the Sanquin FACS facility; G. Stoecklin (University of Heidelberg) for the 4xS1m aptamer construct; T. Schumacher (Netherlands Cancer Institute) for providing MART-1 TCR viral supernatants and Mel526 and Mel888 tumor cell lines; M. Hoogenboezem, N. Zandhuis, S. Castenmiller, N. Sostaric and I. Foskolou for technical help and D. Amsen, M. Nolte and B. van Steensel for critical reading of the manuscript.

This research was supported by Dutch Cancer Society (KWF Kankerbestrijding, 10132), European Research council (ERC) consolidator award PRINTERS 817533 and Oncode Institute, all to M.C.W., and the Amsterdam Infection and Immunity (AII) Postdoc Grant (25262) to B.P.

## Data deposition

The MS data of the RNA pull-down and the co-IP have been deposited to the ProteomeXchange Consortium via the PRIDE partner repository with the dataset identifier PXD028171.

## Notes

### Competing Interest Statement

The authors have declared no competing interest.

### Summary of Updates

Further validation experiments have been included

