## Supplementary figures and images for "Time-dependent regulation of cytokine production by RNA binding proteins defines T cell effector function"

### Figure S1

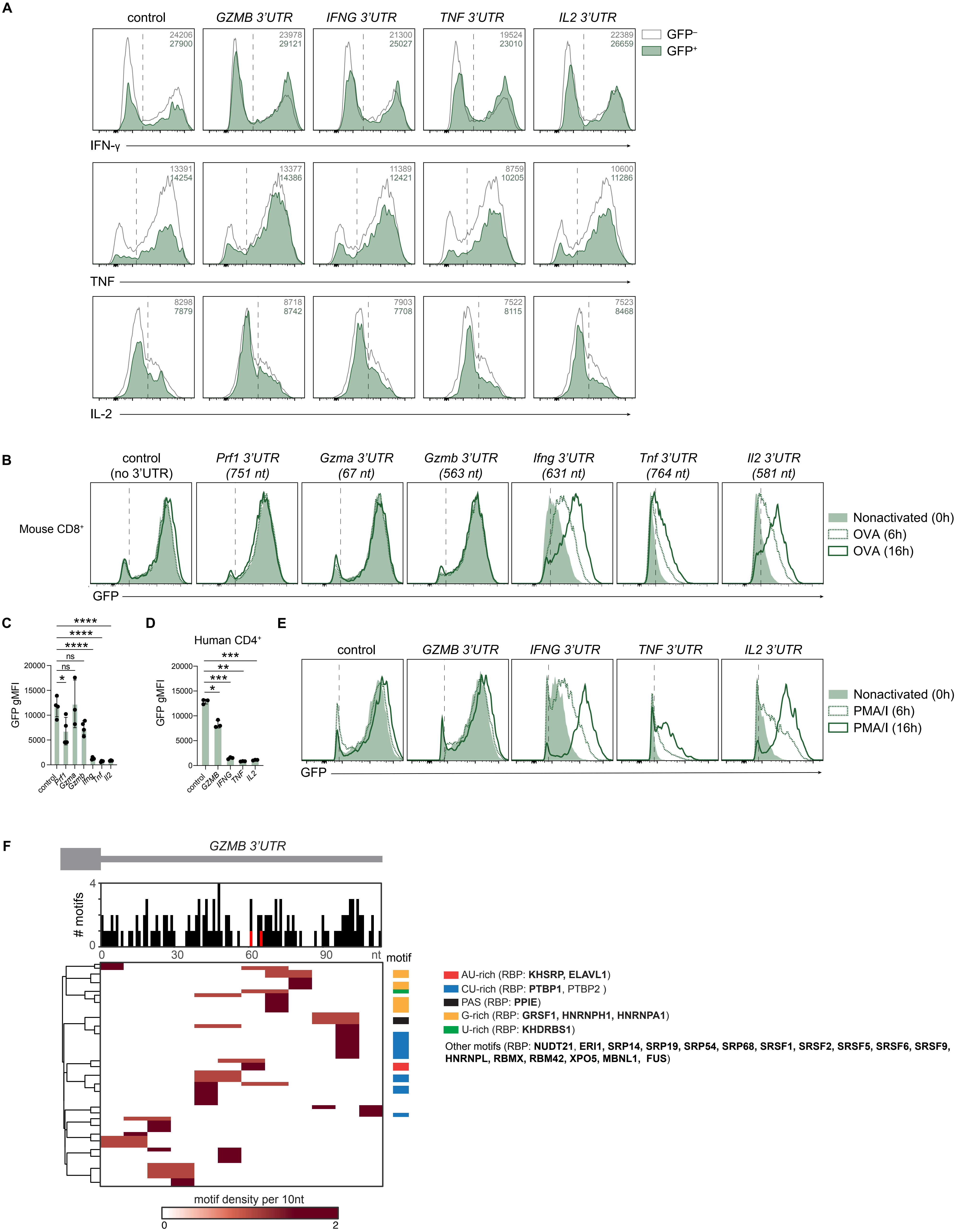

### Figure S2

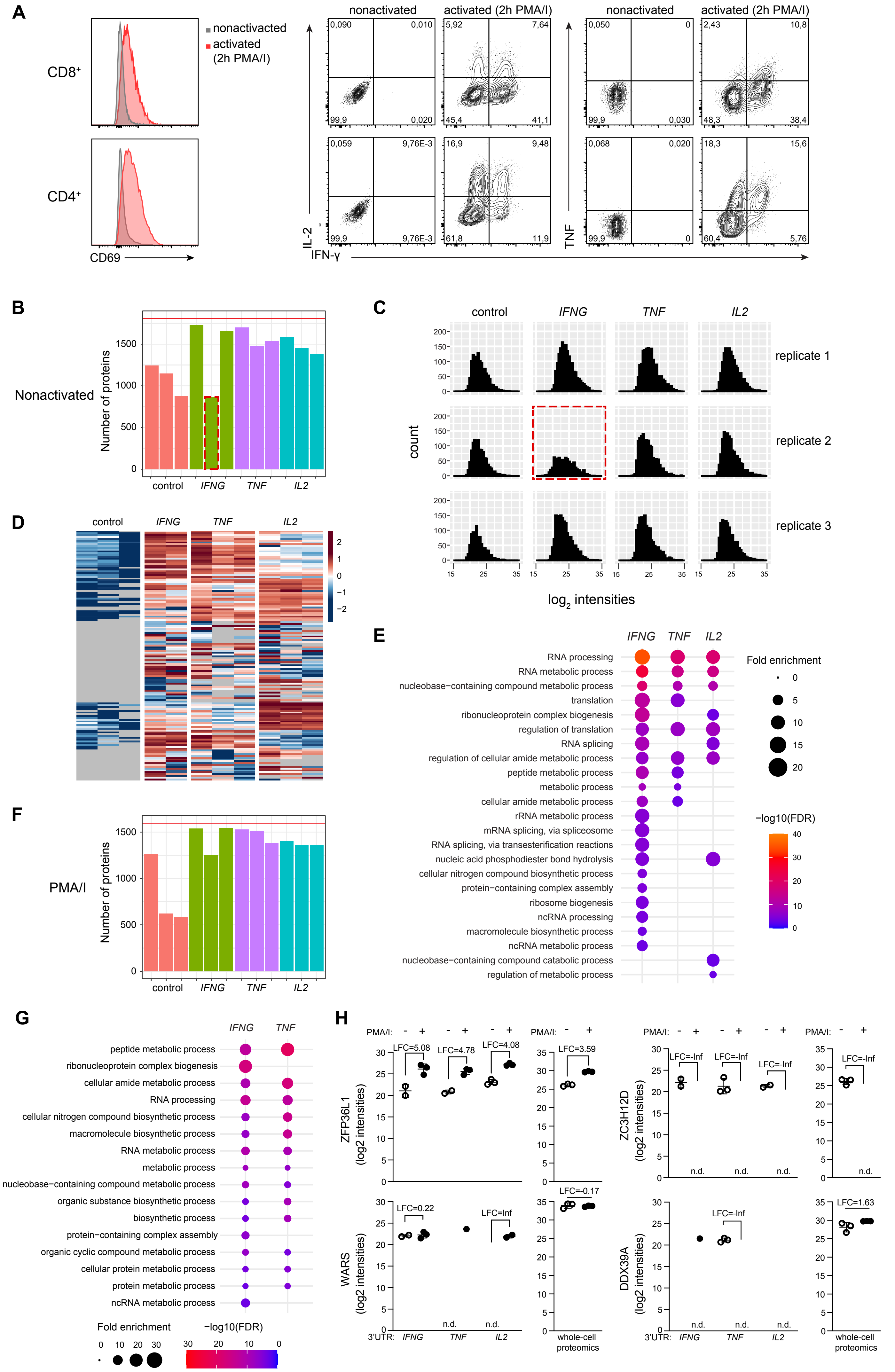

### Figure S5

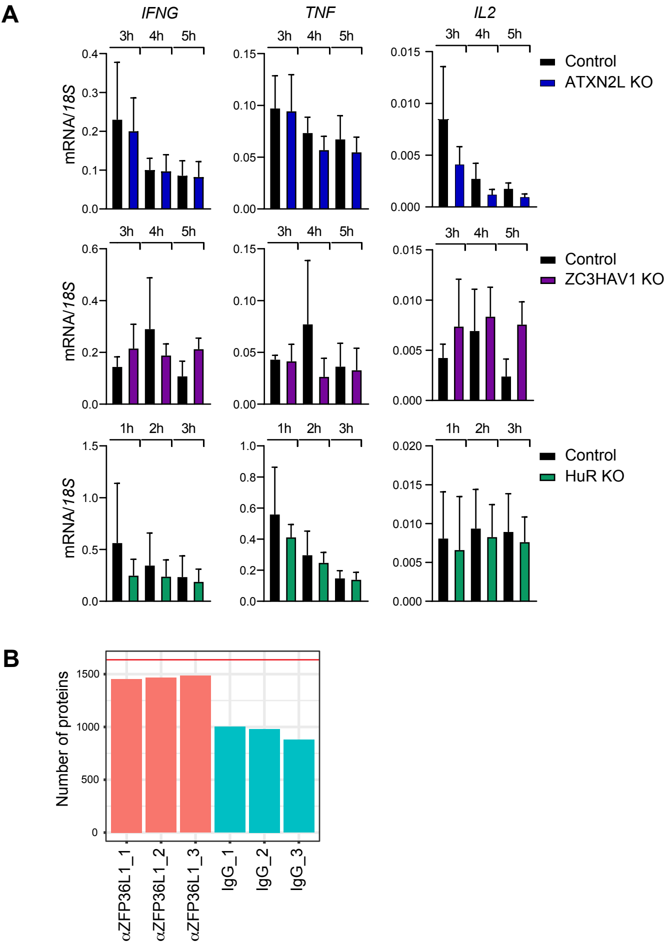

### Figure S6

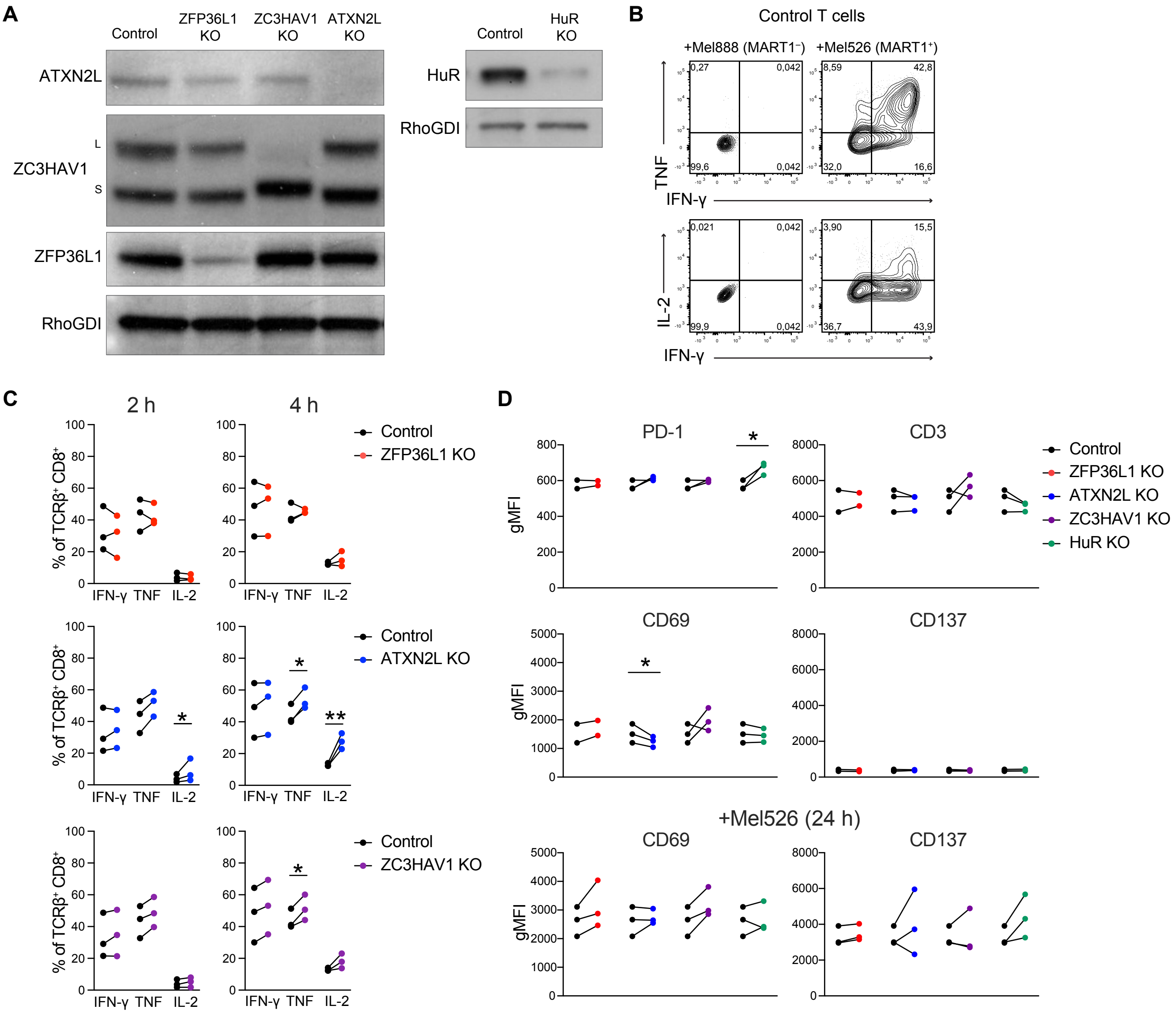

### Figure S7

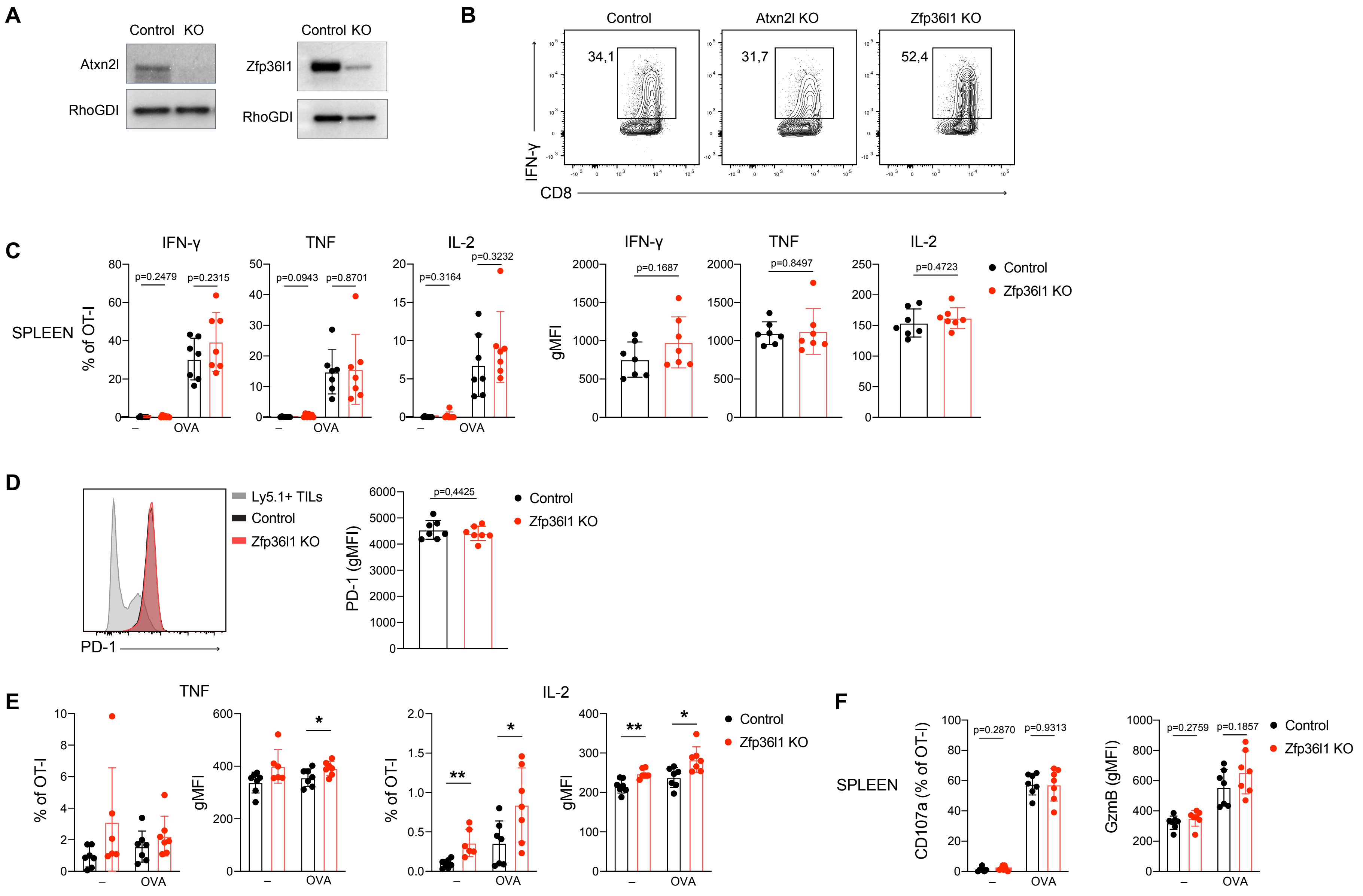
