## Supplementary material for "Time-dependent regulation of cytokine production by RNA binding proteins defines T cell effector function": Figure S3

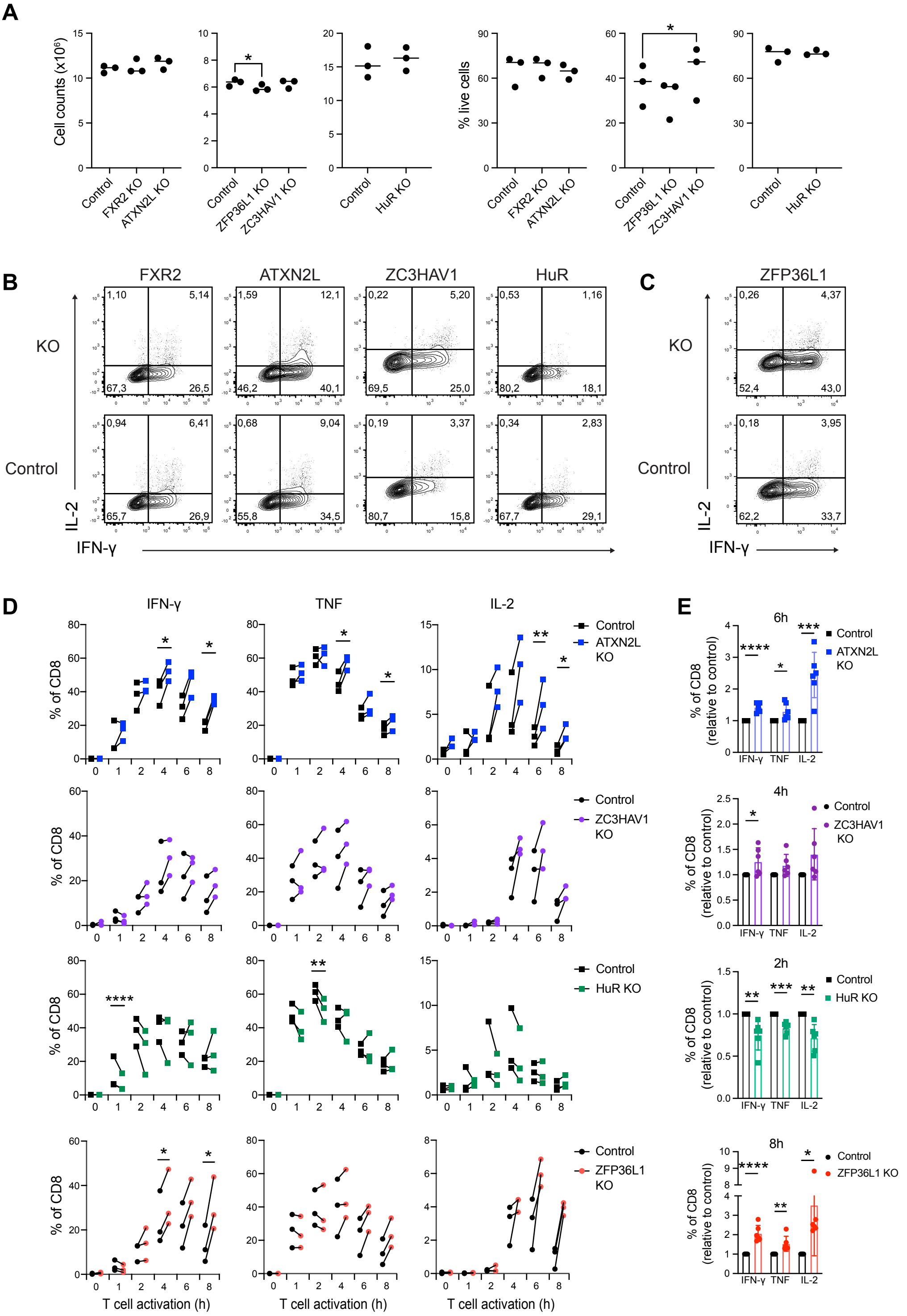


F

| | early kinetics ① | amplitude ② | late kinetics ③ |
|---|---|---|---|
| ATXN2L KO | accelerated | ↑ | - |
| HUR KO | delayed | ↓ | - |
| ZC3HAV1 KO | - | ↑ | - |
| ZFP36L1 KO | - | ↑ | prolonged |
