## Supplementary material for "Time-dependent regulation of cytokine production by RNA binding proteins defines T cell effector function": Figure S4

**A** Human CD4$^+$

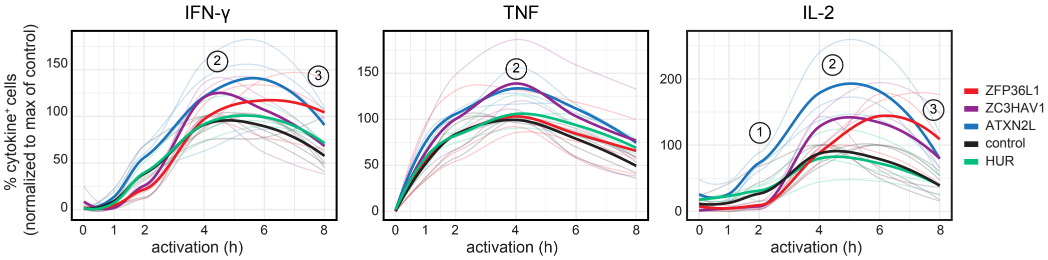


| | early kinetics ① | amplitude ② | late kinetics ③ |
|---|---|---|---|
| ATXN2L KO | accelerated | ↑ | - |
| HUR KO | - | - | - |
| ZC3HAV1 KO | - | ↑ | - |
| ZFP36L1 KO | - | ↑ | prolonged |
