## Supplementary material for "Time-dependent regulation of cytokine production by RNA binding proteins defines T cell effector function": Table S6

### Dataset S6

| Primer name | sequence |
|---|---|
| Murine GZMB 3'UTR F | 5’-GGGGTCGACCTACAGAAGCAACATGGATCC-3’ |
| Murine GZMB 3'UTR R | 5’-CCCTTTGCGGCCGCTTTTATTTGATTTTACATCATTTTTGTCC-3’ |
| Murine IFNG 3'UTR F | 5’-CCATCGATGGATCCGTCGACTGCTGATTCGGGGTGGGG-3’ |
| Murine IFNG 3'UTR R | 5’-CATCGATGCGGCCGCGGTTGCAAAGGTATACTTTATTC-3’ |
| Murine TNF 3'UTR F | 5’-CCATCGATGGATCCGGGAATGGGTGTTCATCCATTC-3’ |
| Murine TNF 3'UTR R | 5’-CCCTTTGCGGCCGCTTTATTTCTCTCAATGACCCGTAGG-3’ |
| Murine IL2 3'UTR F | 5’-CCATCGATGGATCCCTATGTACCTCCTGCTTACAACAC-3’ |
| Murine IL2 3'UTR R | 5’-<br>CCCTTTGCGGCCGCTTTTTTTTTTTTTTTAGAGGAGAGCTTTATTTC-<br>3’ |
| Human GZMB 3’UTR F | 5’-CCCGGATCCCTACAGGAAGCAAACTAAGCCCC-3’ |
| Human GZMB 3’UTR R | 5’-GGGGCGGCCGCTTTTTTTTTTTTTTTTTTTTTTTTTTTTTTTTCCACT<br>CAG-3’ |
| Human IFNG 3’UTR F | 5’-CGGGATCCGGTTGTCCTGCCTGCAATATTTG-3’ |
| Human IFNG 3’UTR R | 5’-CCCTTTGCGGCCGCTTTTTTTTTTTTTTTTTTTTTTTTTTTTTTGTT<br>GTGAACTTACACTTTATTC-3’ |
| Human TNF 3’UTR F | 5’-CCATCGATGGAGGACGAACATCCAACCTTCC-3’ |
| Human TNF 3’UTR R | 5’-GGGGGATCCTTTTTTTTTTTTTCTTTTCTAAGCAAACTTTATTTC<br>TCGCC-3’ |
| Human IL2 3’UTR F | 5’-CCATCGATTAATTAAGTGCTTCCCACTTAAAAC-3’ |
| Human IL2 3’UTR R | 5’-GGGGGATCCTTTTTTTTTTTTTTTTTTTTTTTTTTATATTTATCAA<br>ATTTATTAAATAG-3’ |
| **CRISPR Cas9 crRNA** | **sequence** |
| Human ZFP36L1_1 | 5'-AAACGGTGCCTGTAAGTACG-3' |
| Human ZFP36L1_2 | 5'-GTCTCGCGAGCTCAGAGCGG-3' |
| Human FXR_1 | 5'- GGAGCCGGGACTGCCCGTCG-3' |
| Human FXR_2 | 5'- GGTAGCCGGACATCCCCAAA-3' |
| Human FXR_3 | 5'- TCCCTTCATCATCCGCACCC-3' |
| Human ATXN2L_1 | 5'-AACTTACCACAACAGCTGTA-3' |
| Human ATXN2L_2 | 5'-CTTGAAGATACCCTCATAAG-3' |
| Human ZC3HAV1_1 | 5'-AAAATCCTGTGCGCCCACGG-3' |
| Human ZC3HAV1_2 | 5'-GTCTCTGGCAGTACTTGCGA-3' |
| Human ZC3HAV1_3 | 5'-CAGAGATGCAGGTTATCGCA-3' |
| Human ELAVL1_1 | 5'-TGTGAACTACGTGACCGCGA-3' |
| Murine ZFP36L1_1 | 5'-GAGTGACCGAGTGCCTGCGA-3' |
| Murine ZFP36L1_2 | 5'-GTCTCGCGAGCTCAGAGCGG-3' |
