## Supplementary material for "Time-dependent regulation of cytokine production by RNA binding proteins defines T cell effector function": Table S7

**RBPs$ID**
HRSP12
MRPL2
EIF4G1
TRUB2
STAU2
EWSR1
DDX10
UTP18
PDCD11
RPS3A
KIAA0922
UBE2L3
GTPBP8
RPS14
ESRP2
TUFM
DNM1L
BCLAF1
FUBP3
MALSU1
SFPQ
PSPC1
DSP
SSB
NIN
LRPPRC
RDX
PA2G4
FLNA
RPL3
RBM19
CALML5
PWP2
RPLP0
UPF1
TLN1
MEX3D
BHMT
PHGDH
RPL7A
NSUN5
NUPL2
DHX9
DDX50
ABCE1

RBMX
PSMA6
EDF1
PPID
WDR46
ALDOA
EXOSC4
PUM1
WDR36
LENG9
DNAJB12
PUM2
DHX30
TRNAU1AP
H1FX
RBM14
SYNCRIP
DBNL
TRA2B
TRA2A
HDLBP
TBL2
DDX17
HNRNPUL1
PEBP1
TBL3
FAM195A
HNRNPK
PUF60
TPD52
PTCD2
RSG1
RPL8
PRDX1
HNRNPH2
RPS2
RBM12
PTPRG
CPSF7
MRPL3
FAM207A
SEC62
RPL37A
ZFP36L2
HNRNPA0
YWHAQ

C11orf68
SAFB2
FKBP3
PTCD3
SF1
NHP2L1
RRS1
RPS27A
RPL13A
WDR75
NOP9
BRIX1
RBM28
SNRPF
CSDE1
FUBP1
NONO
DUS3L
SON
SRSF5
DCD
KTN1
SAFB
ALKBH1
FXR1
LSM14A
PRKDC
LARP1
AHCYL2
LARP4
ACAT1
MOV10
UBAP2L
SRSF9
HSPB1
CAPRIN1
RPL5
DDX28
UBE2M
RNH1
ELAVL1
DDX54
SRRT
MATR3
CLTB
ARFGAP3

FBL
RPS27
HPRT1
HNRNPU
FAM120A
DSTYK
PDCD6IP
FASTKD2
SPTAN1
HNRNPR
FXR2
RRP7A
PAK2
DAZAP1
MRPL13
EPRS
DDX52
JMJD1C
ELAC2
UTP15
SND1
URB1
RBM45
RPS12
TUBA4A
RBM47
A1CF
HNRNPA2B1
CTTN
MTERFD1
WBSCR16
YWHAE
HADHB
P4HB
ADAR
RPS11
C7orf25
ARHGEF2
NCL
ILF3
PDIA3
RRP12
TSN
KIAA0020
ZC3H7A
MTDH

HNRNPA3
MRPS5
SNRPD2
SNRNP200
CCDC86
DNMT1
EIF4A1
NOL8
YTHDF1
FUS
ANKRD17
ALDH6A1
LIN28B
PTBP2
PARK7
PRRC2A
CCDC9
ZC3HAV1
HSPD1
SPTBN1
MEX3C
AGO2
HNRNPM
DDX18
DDX5
PRPF40A
ATP5A1
NOP58
RBM33
CHGA
TBRG4
UTP20
XRCC6
NFXL1
PTBP3
PTBP1
TAGLN2
RBPMS
PRPF4B
PNN
SRRM1
MRPL50
HN1L
MRPL28
PES1
USP36

MRPL37
PARD3
EBNA1BP2
EIF3G
EIF3J
AAMDC
VIM
EIF4A2
RRBP1
VCL
ATXN2
MRPS26
SLTM
ST13
HNRNPAB
MSI1
TARDBP
STARD9
GTF2I
DCAF13
SERBP1
RPS4X
EFHD2
U2SURP
PURB
PURA
CIRH1A
CPEB4
RANBP2
TCEB3
NAT10
SARS
GRB2
SAP18
SQSTM1
MTPAP
TMA16
DHX15
RPSA
RPS3
MRPS16
MAT2A
BYSL
PIP
KHDRBS1
YBX1

CSNK1D
CNN3
CSTF3
SNRPE
AHNAK
WDR3
CLINT1
RPS6
EIF3H
RBM10
PABPN1
HNRNPD
HNRNPDL
NAA38
CSRP1
CIRBP
HNRNPA1
DARS
HNRNPH3
PIGR
PPP1R10
HSPE1
SRP54
SNRPA1
ZCCHC7
RBM39
ZCCHC3
SKIV2L2
SRPRB
PARP12
HNRNPLL
SPEN
NOL11
RUVBL1
HNRNPF
RBMS1
GRSF1
ZCCHC9
ZC3H13
MRPS24
SNRPG
CNN1
RPL4
DDX56
RBM15
NCOR2

CTNND1
KIF13B
ANXA11
ILF2
SRP72
RPS15
XRN2
MRPS28
DDX31
TRUB1
EEF2
UBE2O
PRR3
PABPC4
PABPC1
ZC3H11A
XRCC5
MYBBP1A
RPL12
LAMA5
RPL7
HDGF
NACA
RPF2
PPIB
MSI2
MRPL39
NRBP1
DDX21
LARS
KHSRP
IGF2BP1
TPI1
HMGB1
HMGB2
MANF
NOP2
RPLP2
RPL19
MRPS21
ATP5B
PSMC2
RPL17
PTPN12
RPL23
IGF2BP2

IQGAP1
KIF1B
PPIF
HSPA5
NASP
TCEA1
PDAP1
RBM38
MRPL43
EML4
CC2D1A
RPL23A
RPL13
NOM1
AKAP8L
EXOSC2
GTF2F1
EIF4A3
NXF1
PCBP2
PRELP
EIF3A
CELF1
CFL1
S100A9
FARSB
DKC1
POLRMT
SNRPD3
RPS13
RPS25
ANP32B
HSPA4
PRRC2B
TMPO
DEK
EIF5B
TOR4A
RARS
NKRF
PPIL4
LSM14B
PRRC2C
G3BP1
GOLGB1
KRI1

ZC3H14
NSUN2
MRPL18
PREB
DIMT1
GTPBP4
NUSAP1
RPS8
RPL30
TAGLN
STRAP
IMP3
U2AF2
NDRG1
NDRG4
EIF4G2
DIEXF
MRPL27
HNRNPH1
CEBPZ
SNRPN
CCDC137
RBM8A
RBM6
GNL2
ZNF106
TLE3
ANK3
NARS
DLD
RPL18A
TRMT10C
SRSF6
NME1
SF3B4
DHX36
DHX8
SRSF7
RRP8
ERAL1
RPL27A
NOP14
IMP4
METTL16
DDX51
YWHAG

MRPL24
HNRNPUL2
TMCO1
MRPS9
DNAJC1
TRIP6
AZGP1
POLR2A
RPL24
NOP56
SF3A2
RPL6
RBM12B
RBPMS2
CKAP4
TAF15
PCBP1
ABT1
YBX3
SRSF1
MKI67
PPP1R8
MAP2
RPL31
CD3EAP
DTD1
TFAM
CADM1
MAP1B
UTP14A
RSL1D1
ARHGAP11A
HIST1H1E
H2AFY2
RAP1GAP2
EEF1A1
WBP11
CCDC124
SMNDC1
LRRC47
DIP2B
PEG10
CDV3
CSRP2
CUL4B
TFB2M

DDX6
SLC45A4
MTERFD2
NUP35
EIF4B
YBX2
RANBP10
COPB1
EIF3E
FRMD4A
DDX3X
DCN
CRCT1
GPR161
RBMX2
BMS1
C15orf52
RPS19BP1
ARFGEF1
TDRD3
DYNC1I2
SCAMP3
NEMF
USP10
TRMT1L
SH3PXD2B
GRWD1
ALDH2
ARPP19
GSPT1
LSG1
IGF2BP3
PUS1
GNL3L
NUDT21
LAP3
RBM22
TNPO1
AQR
MRPS25
IQCG
SRRM2
HNRNPL
ZNF706
CPSF4
CDC5L

CSTA
GLTSCR2
FTSJ3
PRPF8
RPS24
SEC61B
SAP30BP
SRPK2
TOP3B
RAN
NTPCR
GAR1
HSPA9
RPS28
HSPA6
MRPL4
PAPD5
MDH1
RALY
MACF1
FASN
SYNE1
TNRC6B
RPL35
DYNC1H1
ZCCHC11
EEFSEC
AHCY
KIAA1967
ZC3H18
NOLC1
TARBP2
ETFA
CHD4
RPL21
SF3B1
FAM50A
NSA2
TOM1
MTO1
RBM24
SRSF10
RTF1
RPL38
SPAG9
COL14A1

SPATS2
DHX57
API5
FNDC3B
GNB2L1
TIA1
PHB
MYLK
MBNL1
NOL7
MARCKS
MECP2
MRPS23
TCP1
ZKSCAN1
SEC23IP
HIST1H1B
UBR4
REXO4
MSN
APOBEC3C
AGFG1
SF3A1
CPS1
CAD
KDM1B
SCYL2
LTA4H
MCRS1
TPD52L1
CCT7
NUCKS1
AK2
YWHAH
ZCRB1
CNBP
TIAL1
MCAT
CSTF2T
RPS19
RPUSD4
EEF1G
STIP1
MEPCE
TMEM106B
MYH11

LLPH
MKI67IP
TIMM10
CAMSAP2
EIF4H
NUFIP2
NOC3L
G3BP2
PABPC3
ENSA
GPS1
SLIRP
GEMIN5
LRRC59
SUPT16H
PHLDB1
CHD3
SRBD1
PRPF3
ZC3H15
RSL24D1
DNAJA1
STOML2
VCP
APTX
STMN1
BASP1
ARCN1
SNRPA
RRP1B
PCNP
FMR1
ESF1
WNK1
MRPL19
MRPL1
MRPS11
MRPL45
EIF5
S100A8
UPF2
PFDN2
RPL18
SRSF2
CGN
EXOSC6

CHCHD3
PSMD9
SUB1
SNTB2
CARHSP1
AKAP1
GAB2
USO1
ALDOC
MDH2
RBM34
IMPDH1
ARHGEF12
RPL32
CISD1
FAM98A
MAT1A
CCT3
SARDH
NAA50
TKT
CSE1L
LIN7C
CCT2
GLOD4
ELAVL2
ATXN2L
SRP14
RPL22
LDHB
HADH
HMGB3
RPL36
BTF3
RPN1
PRDX6
DHRS4
RPL29
H2AFY
NOC2L
YTHDF2
CRIPT
RBM4
PDLIM5
MYO5A
MRPS7

GNL3
CTNNB1
SRSF11
CHD8
CDC37
WIBG
STRBP
BAIAP2
ANXA2
SORBS1
PCF11
STIM1
PATL1
RPL14
TNS1
SUPV3L1
NDRG2
TES
RAB1B
HNRNPC
HNRNPCL1
RBM27
ASPM
GOT2
PMS1
NPM1
LPP
CDC42EP4
RPL28
HSP90B1
EIF3D
MRPL32
YARS
CCDC43
NFIC
TESC
QTRT1
ITGA1
CDK11A
KIF5B
EIF3I
CDC42
TMEM214
CWF19L1
LMO7
RBM17

DLG1
PELO
PTCD1
ALYREF
CCDC47
EML1
C11orf58
POP1
SERPINI2
LMNA
UBQLN1
GCN1L1
CORO1C
Sep.09
MRPS31
TECR
CENPF
PACS1
ADD3
UNK
CSTB
CLASP2
HP1BP3
COA6
FRMPD3
PPP1R12C
RAP1GDS1
DYNC1LI1
SMOC1
LRP1
LSM4
IQGAP2
SCG3
PPAN
HCLS1
TMSB4X
PPIA
CYB5A
PFN1
PPP1R12A
ENO1
RBM26
RBM5
FNBP1L
MLLT4
SLK

BRD2
SMARCC2
TMEM230
NUMB
RBMS2
LASP1
MAP4
ZC3H7B
NEB
MARK2
DHX33
IDH1
CNDP2
DNAH2
ITGB1
DST
SH3KBP1
CNOT3
R3HDM1
R3HDM2
IMPDH2
KNOP1
HIST1H4B
TRMT2A
GFM1
DDX46
RPL11
NDC1
GATM
LARP7
RPS5
SRSF3
RPA1
PIN1
MBNL3
FAM120C
MGME1
MTHFD1
MYEF2
CELF2
MFAP1
ARPP21
ESRP1
KHDRBS3
CCAR2
CSTF2

UBAP2
MEX3B
RAVER1
ZNF277
MAP1A
SRSF4
DDX39B
EIF4G3
AKAP8
C11ORF68
TEX10
LARP4B
YTHDF3
SARNP
RBFOX1|RBFOX2
ANKHD1
IREB2
SF3A3
RPS15A
YTHDC1
CHERP
SURF6
CHD2
EIF3CL|EIF3C
DDX39A
PHF5A
TOP1
CPSF6
RBM3
NOL6
TRIM56
DDX1
EFTUD2
RC3H2
TOP2B
ETF1
RBM15B
DDX60L
OTUD4
ZCCHC6
DHX16
SUPT5H
PUS7
RBM25
HNRNPLL|HNRPLL
WDR43

MPHOSPH10
SNRPD1
MKRN2
A0A087WWQ2|EMG1|V9GYP5
TTF1
MRPS18A
LONP1
HELZ
PNPT1
DDX19A
PAK1IP1
PRPF38B
MTHFSD
ESPL1
UTP6
DRG1
GIGYF2
DZIP3
ARL6IP4
ISG15
CNOT1
BZW1
ABCF1
RNPS1
DHX29
POLR2B
YLPM1
CMSS1
RBBP4
CASC3
RTCB
BZW2
RBMXL1
DDX55
UFL1
HEATR1
EIF1AY|EIF1AX
ZNF326
UTP11
C7ORF55|LUC7L2|LUC7L
RNF213
UBTF
EIF2S1
RCC2
KPNA2
THRAP3

ASCC3
HERC5
SDAD1
PRPF31
GPATCH4
FCF1
PPIG
NFX1
DDX42
RPL15
ZNF638
ZFR
SNRPB|SNRPN
SUMO1
SUMO3|SUMO2
NAA15
TOP2A
STAU1
LSM2
AARS
ZNF622
SUPT6H
SRPK1
SMG6
FASTKD1
MYH9
CARM1
SNRPGP15|SNRPG
LDHA
CEPT1
RPL10
ABCF2
KIF1B|KIF1BBETA
CPEB1
PUM3
NCBP3
XPO5
MRM3
UTP4
LAS1L
RNMT
RPL27
MMTAG2
RCL1
C1QBP
SUGP2

DDX27
ZNFX1
UBA1
DHX37
SLC25A5|SLC25A6
PNO1
RACK1
GART
SRSF8
PRMT1
RPL26L1|RPL26|J3QQQ9
NOL9
CCT4
PHF6
FYTTD1
FLNB
CLUH
EIF2AK2
DIAPH1
ZRANB2
CORO1A
AATF
ARF3|ARF1
TCOF1
IFI16
ACIN1
MAP4K4
SNRNP70
TUBB
ALKBH5
POLDIP3
TXN
DDX49
YTHDC2
RBM4B
PKM
SCAF11
EIF2A
FYB
UPF3B
XRN1
CHD1
RRP1
PHRF1
KIF2A
DDX47

EIF3L
DNTTIP2
RPL7L1
TCERG1
MRPS27
PARP1
SECISBP2L
DIS3L2
EXOSC10
TRIP12
AP3D1
LYAR
SRFBP1
SSRP1
NMD3
BK150C2|APOBEC3D
MCM3
GDI2
URB2
ZNF598
NOL10
U2AF1L5|U2AF1
SSBP1
BLMH
EZR
HSP90AB1
LSM8
KRR1
DDX24
TRIM28
ARHGDIB
BOP1
RPS9
PARN
RYDEN
FARSA
EXOSC9
RIDA|HRSP12
FRG1
TRAP1
GP2
HSPA8
LUC7L3
PRDX2
LCP1
WDR1

GAPDH
H3F3A|H3F3B
MORC2
NOA1
RPS7
PGK1
RPL34
NIFK
EEF1A1|EEF1A1P5
RPL10A
DSC1
HIST1H4A
METAP1
NMT1
RPL37
SERPINB12
RPL22L1
RIOK1
UBA52
TGM1
HSD17B10
NOC4L
C9ORF114
DSG1
HSP90AA1
RPS16
IDH2
GSDMA
TRIM25
RSRC2
POF1B
PDIA4
A0A0B4J203|FIP1L1
RPS23
HIST1H2BM
PPRC1
PLA2G1B
RBBP6
SERPINB4|SERPINB3
TUBA1B
KPRP
XP32
JUP
MEX3A
RPL36A
FAU

NGDN
HIST1H1C
H1F0
SERPIND1
ZRSR2
ZRSR1
ZNHIT6
ZNF768
ZNF579
ZNF473
ZNF385A
ZNF346
ZNF239
ZMAT5
ZMAT3
ZMAT2
ZGPAT
ZFR2
ZFP36L1
ZFP36
ZFC3H1
ZCCHC8
ZCCHC5
ZCCHC24
ZCCHC2
ZCCHC17
ZCCHC14
ZCCHC13
ZC3HC1
ZC3HAV1L
ZC3H8
ZC3H6
ZC3H4
ZC3H3
ZC3H12D
ZC3H12C
ZC3H12B
ZC3H12A
ZC3H10
YRDC
YARS2
XPOT
XPO7
XPO6
XPO4
XPO1

XAB2
WRAP53
WDR83
WDR61
WDR5
WDR4
WDR12
WBP4
WARS2
WARS
VARSL
VARS2
VARS
UTP3
UTP23
UTP14C
UTP11L
USP39
USB1
URM1
UPF3A
UNKL
UHMK1
U2AF1L4
U2AF1
TYW5
TYW3
TYW1
TXNL4B
TXNL4A
TUT1
TTF2
TST
TSR3
TSR2
TSR1
TSNAX
TSFM
TSEN54
TSEN34
TSEN2
TSEN15
TRPT1
TROVE2
TRNT1
TRMU

TRMT61B
TRMT61A
TRMT6
TRMT5
TRMT44
TRMT2B
TRMT13
TRMT12
TRMT112
TRMT11
TRMT10B
TRMT10A
TRMT1
TRIT1
TRIM71
TRIM21
TRDMT1
TPR
TOE1
TNRC6C
TNRC6A
TNPO3
TNPO2
TLR8
TLR7
TLR3
TIPARP
THUMPD3
THUMPD2
THUMPD1
THOC7
THOC6
THOC5
THOC3
THOC2
THOC1
THG1L
TGS1
TFIP11
TFB1M
TEX13A
TERT
TEP1
TEFM
TDRKH
TDRD9

TDRD7
TDRD6
TDRD5
TDRD15
TDRD12
TDRD10
TDRD1
TARSL2
TARS2
TARS
TARBP1
TAF9
SYMPK
SYF2
SWT1
SUZ12
SUPT4H1
SUGP1
SSU72
SRSF12
SRRM4
SRRM3
SRPR
SRP9
SRP68
SRP19
SREK1
SRA1
SPATS2L
SNW1
SNUPN
SNRPC
SNRPB2
SNRPB
SNRNP48
SNRNP40
SNRNP35
SNRNP27
SNRNP25
SNIP1
SMN2
SMN1
SMG9
SMG8
SMG7
SMG5

SMG1
SMAD9
SMAD7
SMAD6
SMAD5
SMAD4
SMAD3
SMAD2
SMAD1
SLU7
SLC4A1AP
SLBP
SKIV2L
SIDT2
SIDT1
SHQ1
SFSWAP
SF3B5
SF3B3
SF3B2
SF3B14
SETX
SETD7
SETD1B
SETD1A
SEPSECS
SECISBP2
SCAF8
SCAF4
SCAF1
SBDS
SART3
SART1
SARS2
SAMHD1
SAMD4B
SAMD4A
RUVBL2
RTCA
RSRC1
RRP9
RRP36
RRP15
RRNAD1
RQCD1
RPUSD3

RPUSD2
RPUSD1
RPS4Y2
RPS4Y1
RPS29
RPS27L
RPS26
RPS21
RPS20
RPS18
RPS17L
RPS17
RPS10
RPP40
RPP38
RPP30
RPP25L
RPP25
RPP21
RPP14
RPLP1
RPL9
RPL41
RPL3L
RPL39L
RPL39
RPL36AL
RPL35A
RPL26L1
RPL26
RPL10L
RPF1
RP9
RNPC3
RNMTL1
RNGTT
RNF17
RNF113B
RNF113A
RNASET2
RNASEL
RNASEK
RNASEH2C
RNASEH2B
RNASEH2A
RNASEH1

RNASE9
RNASE8
RNASE7
RNASE6
RNASE4
RNASE3
RNASE2
RNASE13
RNASE12
RNASE11
RNASE10
RNASE1
RIOK3
RIOK2
REXO2
REXO1
REPIN1
RDM1
RC3H1
RBMY1J
RBMY1F
RBMY1E
RBMY1D
RBMY1B
RBMY1A1
RBMXL3
RBMXL2
RBMS3
RBM7
RBM48
RBM46
RBM44
RBM43
RBM42
RBM41
RBM23
RBM20
RBM18
RBM11
RBFOX3
RBFOX2
RBFOX1
RAVER2
RARS2
RANBP6
RANBP17

RALYL
RAE1
R3HCC1L
R3HCC1
QTRTD1
QRSL1
QKI
QARS
PUSL1
PUS7L
PUS3
PUS10
PURG
PTRHD1
PTRH2
PTRH1
PTRF
PTGES3L-AARSD1
PTGES3
PSTK
PSMA1
PSIP1
PRPF6
PRPF40B
PRPF4
PRPF39
PRPF38A
PRPF19
PRPF18
PRKRA
PRIM1
PQBP1
PPWD1
PPIL3
PPIH
PPIE
PPARGC1B
PPARGC1A
POP7
POP5
POP4
POLR2L
POLR2K
POLR2J3
POLR2J2
POLR2J

POLR2I
POLR2H
POLR2G
POLR2F
POLR2E
POLR2D
POLR1E
PNRC2
PNLDC1
PLRG1
PLD6
PIWIL4
PIWIL3
PIWIL2
PIWIL1
PINX1
PIN4
PIH1D3
PIH1D2
PIH1D1
PHAX
PET112
PDE12
PDCD7
PDCD4
PCBP4
PCBP3
PATL2
PARS2
PARP4
PAPOLG
PAPOLB
PAPOLA
PAPD7
PAPD4
PAN3
PAN2
PAIP2B
PAIP2
PAIP1
PABPN1L
PABPC5
PABPC4L
PABPC1L2B
PABPC1L2A
PABPC1L

OBFC1
OASL
OAS3
OAS2
OAS1
NYNRIN
NXT2
NXT1
NXF5
NXF3
NXF2B
NXF2
NUTF2
NUP153
NUFIP1
NUDT16L1
NUDT16
NSUN7
NSUN6
NSUN4
NSUN3
NSRP1
NR0B1
NPM3
NPM2
NOVA2
NOVA1
NOP16
NOP10
NOL3
NOL12
NOB1
NIP7
NHP2
NELFE
NCBP2L
NCBP2
NCBP1
NARS2
NANOS3
NANOS2
NANOS1
NAF1
N4BP1
MVP
MTRF1L

MTRF1
MTIF3
MTIF2
MTG1
MTFMT
MSL3
MRTO4
MRRF
MRPS6
MRPS36
MRPS35
MRPS34
MRPS33
MRPS30
MRPS22
MRPS2
MRPS18C
MRPS18B
MRPS17
MRPS15
MRPS14
MRPS12
MRPS10
MRPL9
MRPL55
MRPL54
MRPL53
MRPL52
MRPL51
MRPL49
MRPL48
MRPL47
MRPL46
MRPL44
MRPL42
MRPL41
MRPL40
MRPL38
MRPL36
MRPL35
MRPL34
MRPL33
MRPL30
MRPL23
MRPL22
MRPL21

MRPL20
MRPL17
MRPL16
MRPL15
MRPL14
MRPL12
MRPL11
MRPL10
MRP63
MRM1
MPHOSPH6
MOV10L1
MKRN3
MKRN1
MIF4GD
METTL5
METTL3
METTL2B
METTL2A
METTL14
METTL10
METTL1
MCTS1
MBNL2
MAZ
MARS2
MARS
MAK16
MAGOHB
MAGOH
MAEL
LUZP4
LUC7L2
LUC7L
LSMD1
LSM7
LSM6
LSM5
LSM3
LSM12
LSM11
LSM10
LSM1
LRRFIP2
LRRFIP1
LIN28A

LCMT2
LARS2
LARP6
LARP1B
L1TD1
KPNB1
KIN
KIAA0430
KIAA0391
KHNYN
KHDRBS2
KHDC1L
KHDC1
KAT8
KARS
JAKMIP1
ISY1
ISG20L2
ISG20
IPO9
IPO8
IPO7
IPO5
IPO4
IPO13
IPO11
INTS9
INTS8
INTS7
INTS6
INTS5
INTS4
INTS3
INTS2
INTS12
INTS10
INTS1
IGHMBP2
IFIT5
IFIT3
IFIT2
IFIT1B
IFIT1
IFIH1
ICT1
IARS2

IARS
HTATSF1
HNRNPA1L2
HINT3
HEXIM2
HEXIM1
HENMT1
HELZ2
HBS1L
HARS2
HARS
HABP4
GUF1
GTPBP3
GTPBP2
GTPBP10
GTPBP1
GTF3A
GSPT2
GPKOW
GPATCH8
GPATCH1
GNL1
GLE1
GFM2
GEMIN8
GEMIN7
GEMIN6
GEMIN4
GEMIN2
GCFC2
GATC
GARS
FTSJ2
FTSJ1
FTO
FRG1B
FIP1L1
FDXACB1
FBXO17
FBLL1
FASTKD5
FASTKD3
FASTK
FARS2
FAM98C

FAM98B
FAM46A
FAM120B
FAM103A1
EZH2
EXOSC8
EXOSC7
EXOSC5
EXOSC3
EXOSC1
EXOG
EXO1
ERN2
ERN1
ERI3
ERI2
ERI1
ENOX2
ENOX1
ENDOV
ENDOU
ENDOG
EMG1
ELAVL4
ELAVL3
ELAC1
EIF6
EIF5AL1
EIF5A2
EIF5A
EIF4ENIF1
EIF4E3
EIF4E2
EIF4E1B
EIF4E
EIF3M
EIF3K
EIF3CL
EIF3C
EIF3B
EIF2S3L
EIF2S3
EIF2S2
EIF2D
EIF2B5
EIF2B4

EIF2B3
EIF2B2
EIF2B1
EIF2AK4
EIF2AK3
EIF2AK1
EIF1B
EIF1AY
EIF1AX
EIF1AD
EIF1
EFTUD1
EEF2K
EEF1E1
EEF1D
EEF1B2
EEF1A2
EED
EDC4
EDC3
EARS2
DZIP1L
DZIP1
DYNLL1
DXO
DUSP11
DUS4L
DUS2
DUS1L
DROSHA
DRG2
DQX1
DNMT3B
DND1
DNAJC21
DNAJC17
DNAAF2
DIS3L
DIS3
DICER1
DHX58
DHX40
DHX38
DHX35
DHX34
DHX32

DGCR8
DGCR14
DENR
DDX60
DDX59
DDX58
DDX53
DDX43
DDX41
DDX4
DDX3Y
DDX26B
DDX25
DDX23
DDX20
DDX19B
DCPS
DCP2
DCP1B
DCP1A
DBR1
DAZL
DAZ4
DAZ3
DAZ2
DAZ1
DARS2
DAP3
DALRD3
CXorf23
CWF19L2
CWC27
CWC25
CWC22
CWC15
CTU2
CTU1
CTIF
CSTF1
CSDC2
CRYZ
CRNKL1
CPSF4L
CPSF3L
CPSF3
CPSF2

CPSF1
CPEB3
CPEB2
CNP
CNOT8
CNOT7
CNOT6L
CNOT6
CNOT4
CNOT2
CNOT11
CNOT10
CMTR2
CMTR1
CLP1
CLK4
CLK3
CLK2
CLK1
CLASRP
CHTOP
CELF6
CELF5
CELF4
CELF3
CDK9
CDK5RAP1
CDC40
CD2BP2
CCRN4L
CCNT2
CCNT1
CCDC59
CCAR1
CARS2
CARS
CAPRIN2
CANX
CALR3
CALR
CACTIN
C9orf129
C9orf114
C2orf15
C1D
C17orf85

C12orf65
BUD13
BRCA1
BOLL
BICC1
BCDIN3D
BAZ2B
BAZ2A
BARD1
AUH
ATXN1L
ATXN1
ASH1L
ASCC1
ARHGEF28
APOBEC4
APOBEC3H
APOBEC3G
APOBEC3F
APOBEC2
APOBEC1
APEX1
ANGEL2
ANGEL1
ANG
ALKBH8
AKAP17A
AIMP2
AIMP1
AGO4
AGO3
AGO1
AFF4
AFF3
AFF2
AFF1
AEN
ADAT3
ADAT2
ADAT1
ADARB2
ADARB1
ADAD2
ADAD1
ACO1
AC004381.6

AARSD1
AARS2
AAR2
